# Cortex-wide neural interfacing via transparent polymer skulls

**DOI:** 10.1101/387142

**Authors:** Leila Ghanbari, Russell E. Carter, Matthew L. Rynes, Judith Dominguez, Gang Chen, Anant Naik, Jia Hu, Md Abdul Kader Sagar, Lenora Haltom, Nahom Mossazghi, Madelyn M. Gray, Sarah L. West, Kevin W. Eliceiri, Timothy J. Ebner, Suhasa B. Kodandaramaiah

## Abstract

Neural computations occurring simultaneously in multiple cerebral cortical regions are critical for mediating cognition, perception and sensorimotor behaviors. Enormous progress has been made in understanding how neural activity in specific cortical regions contributes to behavior. However, there is a lack of tools that allow simultaneous monitoring and perturbing neural activity from multiple cortical regions. To fill this need, we have engineered “See-Shells” – digitally designed, morphologically realistic, transparent polymer skulls that allow long-term (>200 days) optical access to 45 mm^2^ of the dorsal cerebral cortex in the mouse. We demonstrate the ability to perform mesoscopic imaging, as well as cellular and subcellular resolution two-photon imaging of neural structures up to 600 µm through the See-Shells. See-Shells implanted on transgenic mice expressing genetically encoded calcium (Ca^2+^) indicators allow tracking of neural activities from multiple, non-contiguous regions spread across millimeters of the cortex. Further, neural probes can access the brain through perforated See-Shells, either for perturbing or recording neural activity from localized brain regions simultaneously with whole cortex imaging. As See-Shells can be constructed using readily available desktop fabrication tools and modified to fit a range of skull geometries, they provide a powerful tool for investigating brain structure and function.

## INTRODUCTION

The mammalian cerebral cortex mediates learned and adaptive forms of sensory-motor behaviors and the evolutionary expansion of the cortex underlies many advanced cognitive capabilities in human and non-human primates. Neuroscientists have taken advantage of the modular organization and segregation of the cortex into anatomically and functionally distinct regions and have made enormous progress understanding how computations performed in specific cortical regions engage in behavior. However, operation of the brain cannot be understood only by analysis of its components in isolation. The mechanisms by which neural activity is coordinated across the cerebral cortex to produce a unitary behavioral output are not well understood. Even simple sensory or motor tasks involve processing of information in multiple cortical areas. For example, deflection of a single whisker results in activation distributed across the sensorimotor cortices^1^, locomotion modulates the neural responses in the primary visual cortex with cell-type specificity^2^, and arousal exerts markedly different effects across the cerebral cortex, both spatially and temporally^3^. Further, such long-range information flow is dependent on the internal brain state as well as information learned from past experiences.

Understanding these large-scale computations requires the ability to monitor and perturb neural activity across large regions of the cortex. Until recently, the study of mesoscopic and macroscopic brain networks has been limited to functional magnetic resonance imaging (fMRI) and magnetoencephalography (MEG). However, fMRI and MEG are limited by their spatial and temporal resolution. Two-photon (2P) imaging^4^ has rapidly emerged as a imaging tool of choice for *in vivo* imaging in rodent models due to improved deep imaging over one-photon (1P) methods^5^. The development of optical sensing tools in recent years allow 2P-based cellular resolution monitoring and manipulations in local circuits. Genetically encoded Ca^2+^ indicators (such as GCaMP6) have enabled *in vivo* high-resolution monitoring of activities of hundreds to thousands of neurons^6-11^. The development of red-shifted variants of Ca^2+^ indicators and optogenetic tools open deep cortical regions for optical sensing and perturbation^12-14^. The advent of streamlined strategies to rapidly generate transgenic mice expressing optical reporters has been matched by the recent development of wide-field 2P imaging approaches^15-17^.

Deploying these new optical tools and instrumentation to image large areas of the cerebral cortex requires replacing the overlying opaque skull with a transparent substrate. A widely-used method to achieve chronic optical access to the brain surface involves implanting “optical windows” or “cranial windows” in which sections of the skull are excised and replaced with glass coverslips^18^. To image even larger regions, strategies for refractive index matching^19,20^ and thinned skull preparations^21^ have been used. These techniques, however, do not reliably allow cellular resolution imaging as image quality is dependent on the surgical preparation and is susceptible to bone regrowth over time. Recently, curved glass windows and associated surgical implantation methodology were introduced that allow chronic optical access to the whole dorsal cortex with cellular resolution imaging^22^. While each of these approaches has advanced the field, each has limitations. Ideally, wide-field optical imaging would be combined with modalities that allow simultaneous perturbation of neural activities to reveal the effect of various brain regions on global cortical activity. Further, combining wide-field optical imaging with simultaneous electrophysiological recordings from different brain regions will provide a better understanding of how global activity patterns modulate activity in local circuits^23^. Therefore, large optical windows with excellent optical properties, long-term functionality,design flexibility, easy fabrication and surgical implantation, and accommodation of other modalities are needed.

Here, we introduce “See-Shells”, digitally designed and morphologically realistic transparent polymer skulls that can be chronically implanted for long durations (>200 days) and allow optical access to 45 mm^2^ of the dorsal cerebral cortex. See-Shells can be customized to fit a variety of skull morphologies, and allow for sub-cellular resolution structural imaging. Further, Ca^2+^ imaging can be performed at both mesoscale and cellular resolution from populations of neurons spread across millimeters of the cortex during awake, head-fixed behavior. See-Shells are easily adapted to include perforations for penetrating stimulation or recording probes. We also demonstrate the ability to perform wide-field Ca^2+^ imaging simultaneously with intracortical microstimulation and extracellular recordings. See-Shells can be inexpensively fabricated using desktop prototyping tools and can be implanted using methodologies adapted from standard cranial window implantation procedures.

## RESULTS

### Device design and fabrication

The overall design of the See-Shells is shown in **Figure 1a**. A motorized stereotaxic instrument was used to profile the skull surface covering the dorsal cortex of an 8-week old C57BL/6 mouse at 85 points. These 85 coordinates provided a point cloud representation of the skull surface used to interpolate a 3D surface that accurately mimicked the skull morphology. This interpolated surface served as a template to digitally design a structural frame for transparent skulls (“See-Shells”) using computer-aided design (CAD) software. The frame was 3D-printed out of polymethylmethacrylate (PMMA) onto which a thin, flexible and transparent polyethylene terephthalate (PET) film was bonded (**Fig. 1a** and **Supplementary Figs. 1-3**). The 3D-printed frame also incorporated screw holes for fastening a custom designed titanium head-plate for head-fixing the animal during experiments.

**Figure 1.**
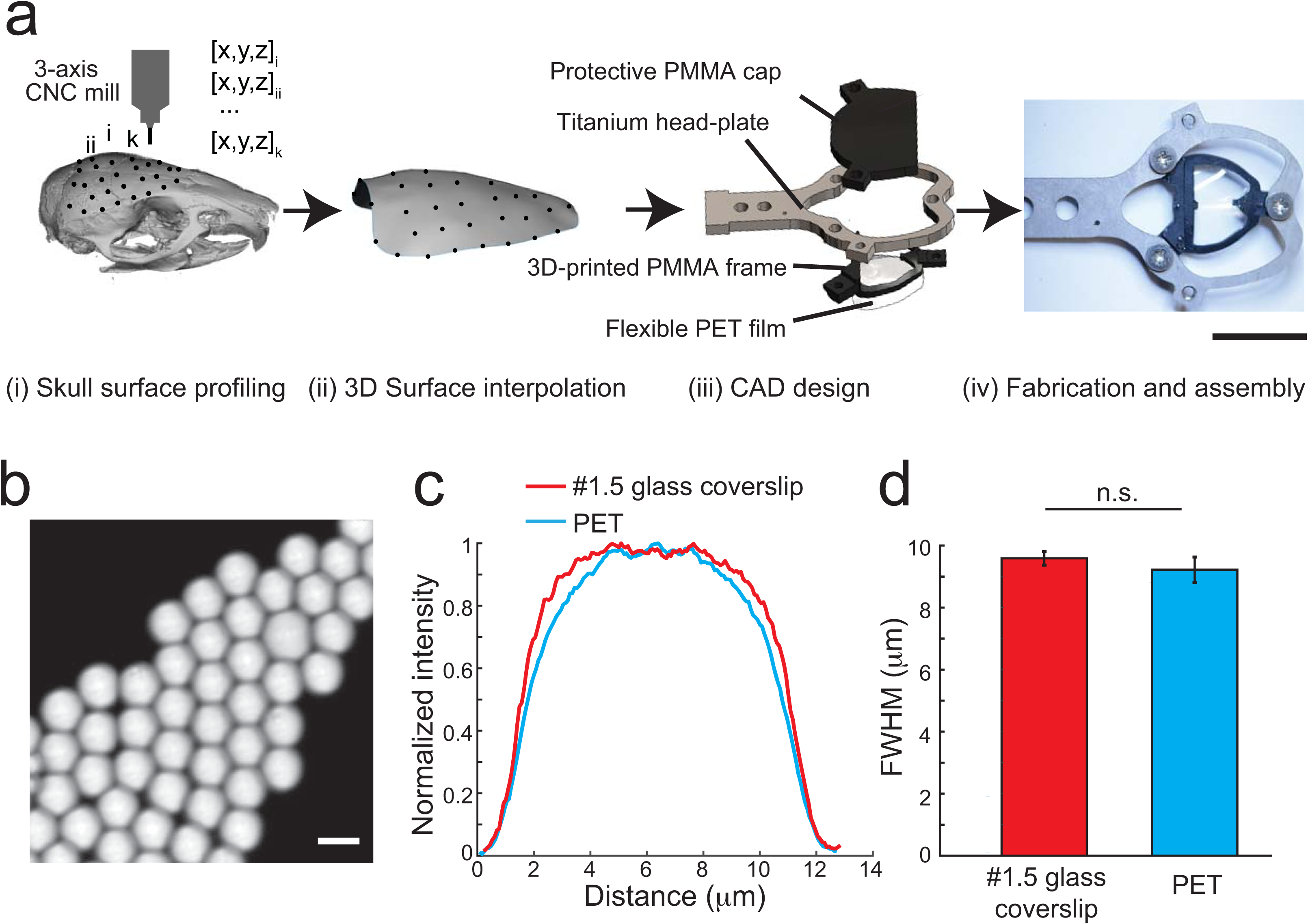
Digitally generating See-Shells: **(a)** (i) Dorsal surface of the mouse skull profiled using a CNC mill integrated into the stereotaxic instrument. (ii) This skull surface profile is used to interpolate a 3D surface. (iii) The 3D surface is used as a template to design morphologically conformant transparent implants (See-Shells), consisting of a 3D-printed Polymethylmethacrylate (PMMA) frame, onto which a thin, optically clear and flexible polyethylene terephthalate (PET) film is bonded. A titanium head-plate fastened to the frame provides mechanical support for head-fixation and a 3D-printed cap protects the implant and underlying brain tissue. (iv) Photograph of a fully fabricated and assembled See-Shell. Scale bar indicates 1 cm. **(b)** 10 µm YG fluorescent beads imaged with 10x imaging objective through the PET film. Scale bar indicates 10 µm. **(c)** Fluorescent intensity profiles measured across the YG beads imaged through PET film and #1.5 glass coverslip. **(d)** Bar plot of full width at half maximum (FWHM) of the intensity profiles of beads imaged through PET film and #1.5 glass coverslip (n = 5 measurements in three PET film each, and 15 measurements in glass coverslip).

PET was chosen for the transparent element as this polymer has excellent optical properties^24^ and is biocompatible^25^. The optical properties of the PET film were compared to the current gold standard, 170 µm thick glass coverslips (#1.5) used for a variety of microscopic imaging experiments. Imaging 10 µm fluorescent beads, which are the size of neural soma, with the PET film and glass coverslips (**Fig. 1b**) revealed no significant difference in the full width at half maximum (FWHM) fluorescence intensity (**Fig. 1c** and **d**, n = 15, t-test, p = 0.10). Additional experiments characterized the light transmittance of three PET samples using a widely tunable laser at common wavelengths used for 2P imaging. All three PET samples yielded light transmittance of more than 90% (**Supplementary Table 1**). On average, across all wavelengths, light transmittance was 91.17 ± 0.94%, 90.85 ± 1.1% and 91.71 ± 0.78% in the three samples tested, compared to 92.79 ± 0.82% in a standard glass coverslip.

Multiphoton and time-correlated single-photon counting (TCSPC) based fluorescence-lifetime imaging (FLIM)^26^ of fluorescent yellow-green (YG) beads was performed to assess whether the PET film introduces changes in light intensity or fluorescence lifetime, respectively. While the present application for See-Shells is fluorescence intensity measurements, FLIM has an additional utility due its sensitivity to changes in the tissue microenvironment and sample conditions without being affected by changes in fluorophore concentration. Concerning light intensity, 2P imaging of YG beads through the PET film required 1-2% higher gain settings on the photomultiplier tube (PMT) compared to glass coverslips. This was expected, as PET film has a slightly lower light transmission efficiency. Given that the imaging was performed well within the PMT specifications, the reduction in light transmission can be easily overcome using a combination of increased laser power and PMT gain settings. Next FLIM imaging was performed using the same 2P instrument. Mean fluorescence-lifetime of three PET film samples were 2.13 ± 0.01 ns, 2.16 ± 0.07 ns, and 2.15 ± 0.02 ns, comparable to the mean fluorescence-lifetime of #1.5 glass coverslips (2.15 ± 0.01 ns). The mean lifetimes are within the resolution limit of this 2P based FLIM system. Therefore, PET has negligible effects on both 2P and FLIM imaging.

### See-Shells can be chronically implanted with no neuroinflammatory effects

See-Shells were chronically implanted on wild-type C57BL/6 (n = 9), Thy1-GCaMP6f (n = 20), and Thy1-YFP mice (n = 3). No bone regrowth was observed in any of the mice and the implants remained clear for several weeks after implantation, with the longest duration assessed at 36 weeks (**Fig. 2a**). As Thy1-GCaMP6f and Thy1-YFP mice are derived from C57BL/6 lines, implants based on the surface profile from the C57BL/6 mouse readily fit these transgenic mice. The digital design methodology used to generate the See-Shells allows easy modifications to fit skull morphologies different from commonly used wild-type mouse strains. As an example, See-Shells were custom designed for the tottering (*tg/tg*) mouse^38^, a strain that has a mutation in the *Cacna1a* gene^27,28^and has narrower skulls than C57BL/6 mice of the same age (**Supplementary Fig. 2**). Similar to chronic implantations on C57BL/6 and derivative mouse strains, the modified See-Shells on *tg/tg* mice (n = 5) remain optically functional for up to 30 weeks (**Fig. 2b**).

**Figure 2:**
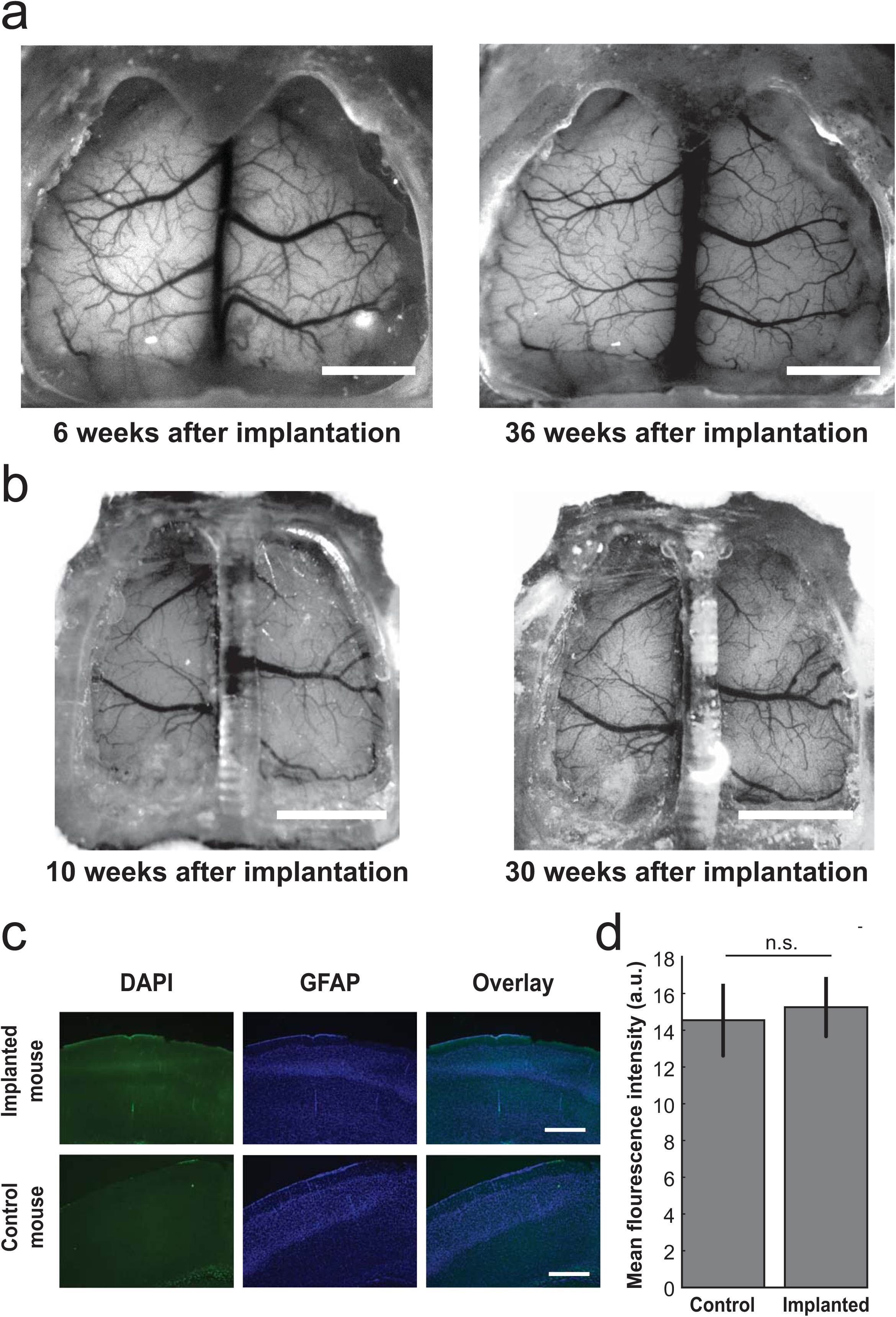
Chronic implantation of See-Shells: **(a)** Photographs of a GCaMP6f mouse 6 and 36 weeks after implantation with See-Shells. Scale bars indicate 2 mm. **(b)** The design can be modified to fit different skull morphologies. Photographs of a *tg/tg* mouse 10 and 30 weeks after implantation. Scale bar indicates 2 mm. **(c)** Immunohistology analysis of mice chronically implanted with See-Shells for 5 weeks compared to age-matched non-surgical control mice. Coronal slices were immunolabeled with anti-glial fibrillary acidic protein (anti-GFAP) and DAPI. No activated astrocytes were observed in any of the mice assessed. Scale bar indicates 500 µm. **(d)** Plot of mean GFAP fluorescence measured in 21 regions of interest (ROIs) in See-Shell implanted and non-surgical control mice.

In a subset of mice (n = 3), the inflammatory effect of chronic implantation was assessed after 5 weeks of implantation by immunostaining for increased expression of glial fibrillary acidic protein (GFAP), a marker for chronic inflammation. No significant increase of GFAP staining was observed in the implanted mice compared to naive control mice (15.26 ±1.64 a.u., n = 21 measurements from 3 mice vs. 14.55 ± 1.99 a.u., n = 21 measurements in 3 mice, t-test, p = 0.2971, **Fig. 2c** and **d**). Thus, the See-Shells can be implanted on mice for long durations of time and allow longitudinal imaging of the dorsal cortex.

### See-Shells allow sub-cellular resolution structural imaging across millimeters of the cortex

Chronically implanted cranial glass windows have been used for high-resolution imaging of neural structure *in vivo* over extended periods of time^29,30^. Several of these studies have revealed key cellular and structural mechanisms underlying experience-dependent plasticity^31-35^. Therefore, we assessed the structural imaging capability of See-Shells across the large field-of-view (FOV) with spatial resolution and imaging depths comparable with glass cranial windows. See-Shells were implanted on Thy1-YFP mice (n = 3) that express the yellow fluorescent protein (YFP) in layer 2/3 and layer 5 pyramidal neurons of the cortex^36^. **Figure 3a** shows a mesoscale 1P image obtained using a wide-field epi-fluorescence microscope at 1x magnification. Multiple locations distributed across the cortex of the same mouse were then imaged at high-resolution with a 2P microscope. Whole cortical columns (360 μm x 360 μm) were reconstructed up to depths of 600 μm (**Fig. 3b).** Capturing multiple z-stacks from adjacent tiles allowed reconstruction of large contiguous volumes of tissue spread across millimeters of the cortex (**Fig. 3c** and **d**). At an optical zoom of 4.4x using a 25x objective, individual neurons and their processes were imaged at depths of ∼300 μm from the pial surface (**Fig. 3e**). At an optical zoom of 7x, finer sub-cellular structures including dendrites and dendritic spines were clearly visible (**Fig. 3f** and **g**). Thus, See-Shells allow the study of fine sub-cellular structures of neurons across the cerebral cortex.

**Figure 3:**
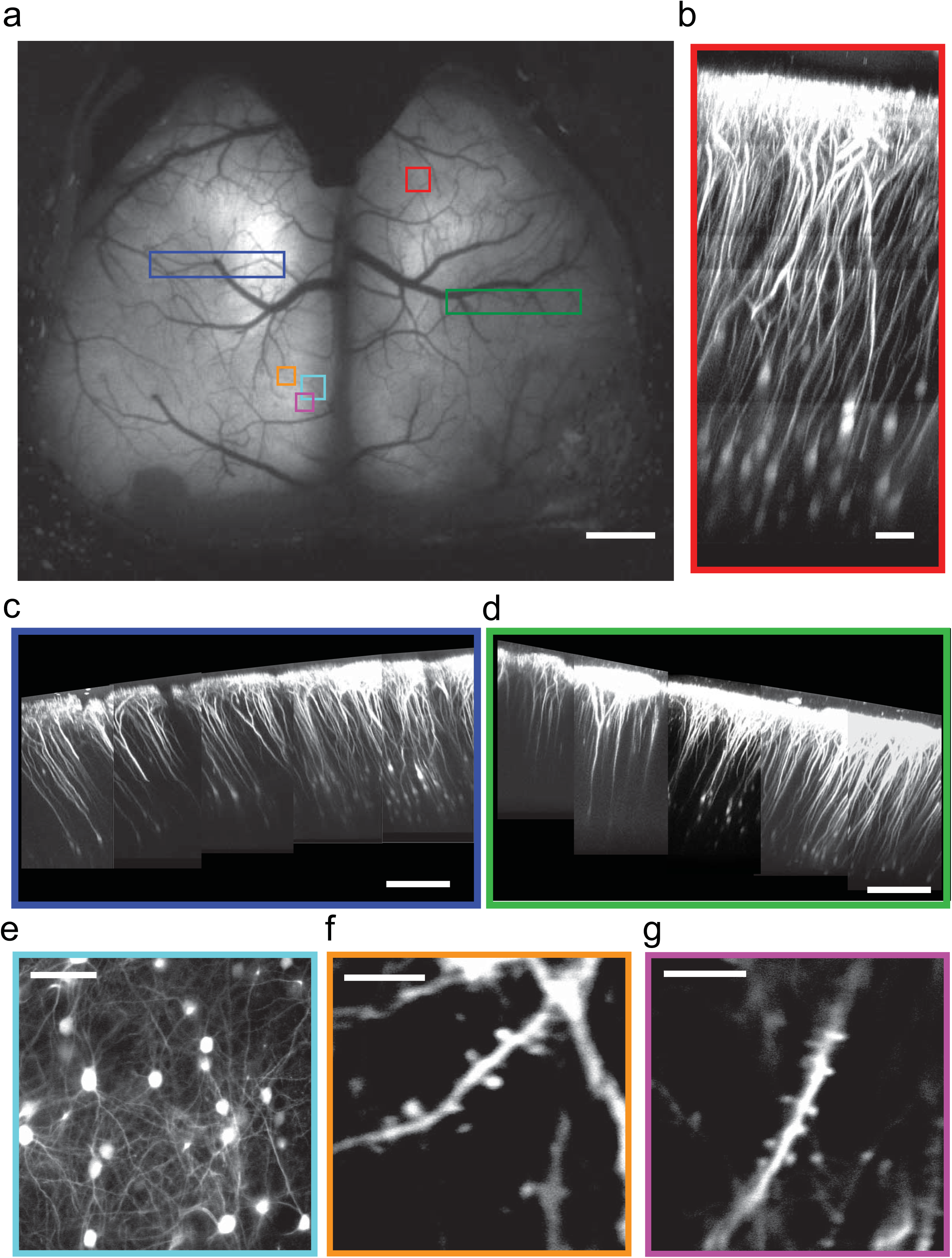
See-shells allow structural imaging from mesoscale to microscale: **(a)** Wide-field image of a Thy1-YFP mouse implanted with a See-Shell taken 2 weeks after implantation. Locations marked with colored blocks were targeted for 2P imaging. Scale bar indicates 1 mm. **(b)** Imaging of a whole cortical column at the location marked by red block in **(a).** Scale bar indicates 80 µm. **(c)** and **(d)** Composite images of cortical columns imaged from large contiguous areas denoted by the blue and green blocks in **(a)**. Scale bar indicates 200 µm. **(e)** High-resolution image of layer 2/3 pyramidal neurons imaged from the cyan block denoted in **(a).** Scale bar indicates 80 µm. **(f)** and **(g)** Dendrites with dendritic spines of layer 2/3 neurons imaged at ∼245 µm depth from the orange and magenta blocks shown in **(a)**. Scale bars indicate 10 µm.

### See-Shells allow Ca^2+^ imaging at multiple spatial scales across the cortex during head-fixed behaviors

To assess the capabilities to monitor cortical neural activity, See-Shells were implanted on Thy1-GCaMP6f mice that express the Ca^2+^ indicator GCaMP6f in layer 2/3 and layer 5 pyramidal neurons^37^. Wide-field imaging using a standard epi-fluorescence microscope captured mesoscale activity across the entire FOV (**Supplementary Fig. 4**, **Supplementary Video 1**). Robust activation of the entire cortex was observed during locomotion, with spontaneous activity observed even at rest. In the mouse shown in **Figure 4a**, four random areas highlighted by the colored blocks were targeted for 2P imaging. At each area, z-stacks of 365 μm x 365 μm were captured when the mouse was fully awake and head-fixed on a custom-built disk treadmill (**Fig. 4b**). Two-dimensional (2D) maximum intensity projections of the z-stacks revealed macroscopic anatomical features such as blood vessels that could be matched with wide-field images to determine the imaging locations *post hoc*. In each tile, time series of Ca^2+^ activity were acquired in planes at 200-300 µm from the pial surface. Individual cells were readily visualized in the average intensity projections of the time series (**Fig. 4c**). Spontaneous Ca^2+^ activity traces from a small subset of the detected neurons in each tile during awake head-fixation are shown in **Figure 4d**.

**Figure 4:**
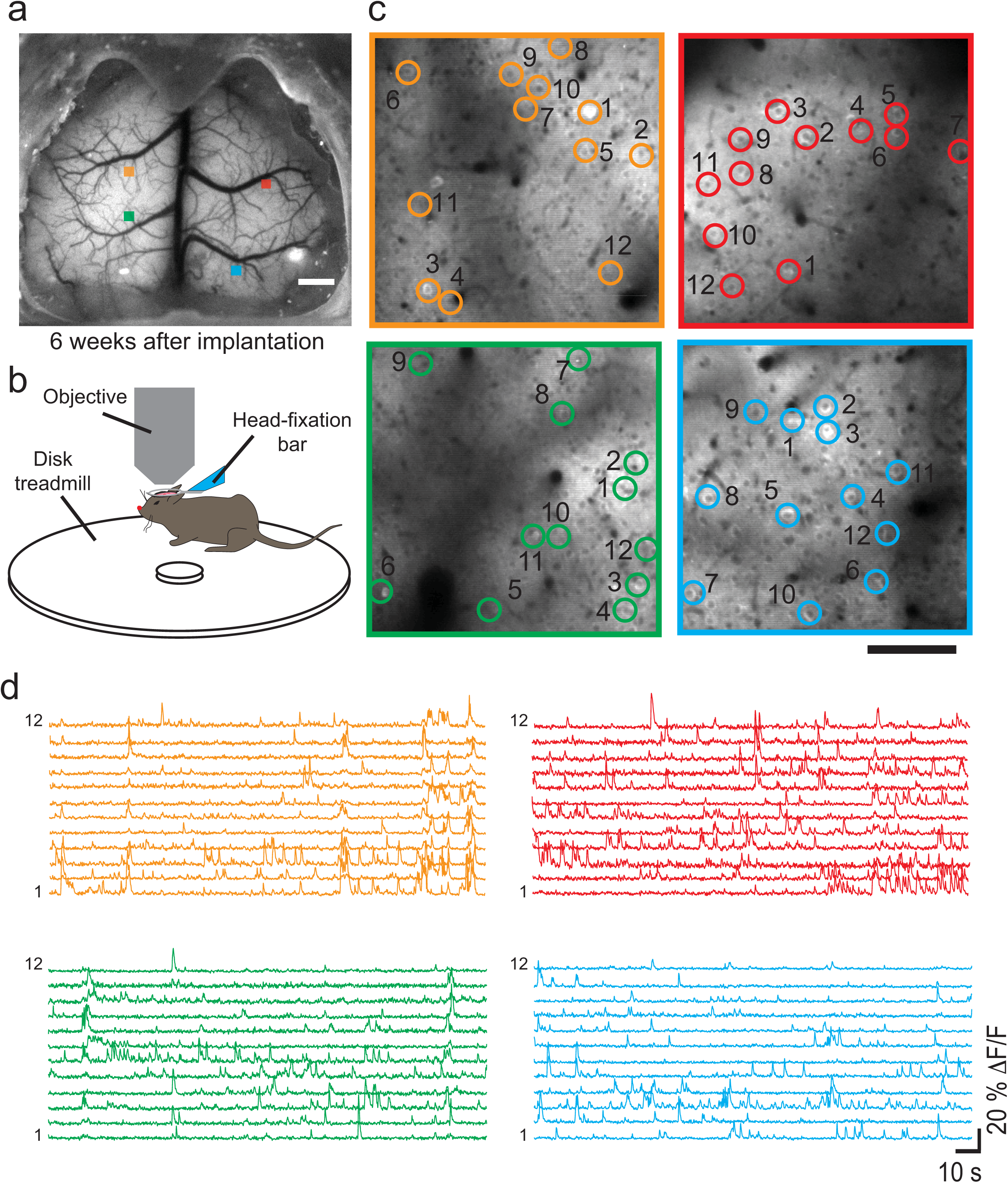
Monitoring Ca^2+^ activities across millimeters of the cortex in awake head-fixed mice: **(a)** Wide-field image of a Thy1-GCaMP6f mouse implanted with a See-Shell taken 6 weeks after implantation. Four locations indicated by the colored blocks were imaged using a 2P microscope. Scale bar indicates 1 mm. **(b)** Schematic of the mouse on the custom design disk treadmill used for 2P imaging. **(c)** Average intensity images calculated from 5-minute time series acquired 200-300 µm deep from pial surface. Scale bar indicates 100 µm. **(d)** Color coded time series of Ca^2+^ activities of neurons identified and annotated by open circles in the respective average intensity images in **(c)**.

Further, Ca^2+^ activities were tracked in Thy1-GCaMP6f mice using 2P imaging in the hindlimb area of the primary motor cortex along with high-speed video monitoring. Mice implanted with See-Shells readily performed a variety of behaviors including walking and grooming during head-fixation. In the recording highlighted in **Supplementary Video 1**, increased Ca^2+^ activity occurred during walking as tracked by movements of the hindlimb, forelimb, and disk treadmill but the modulation was absent during grooming, indicated by forelimb movement (**Supplementary Fig. 5a**). Cellular resolution imaging could be performed in the same mouse for multiple experimental sessions spread across several months (**Supplementary Fig. 5b**).

**Figure 5:**
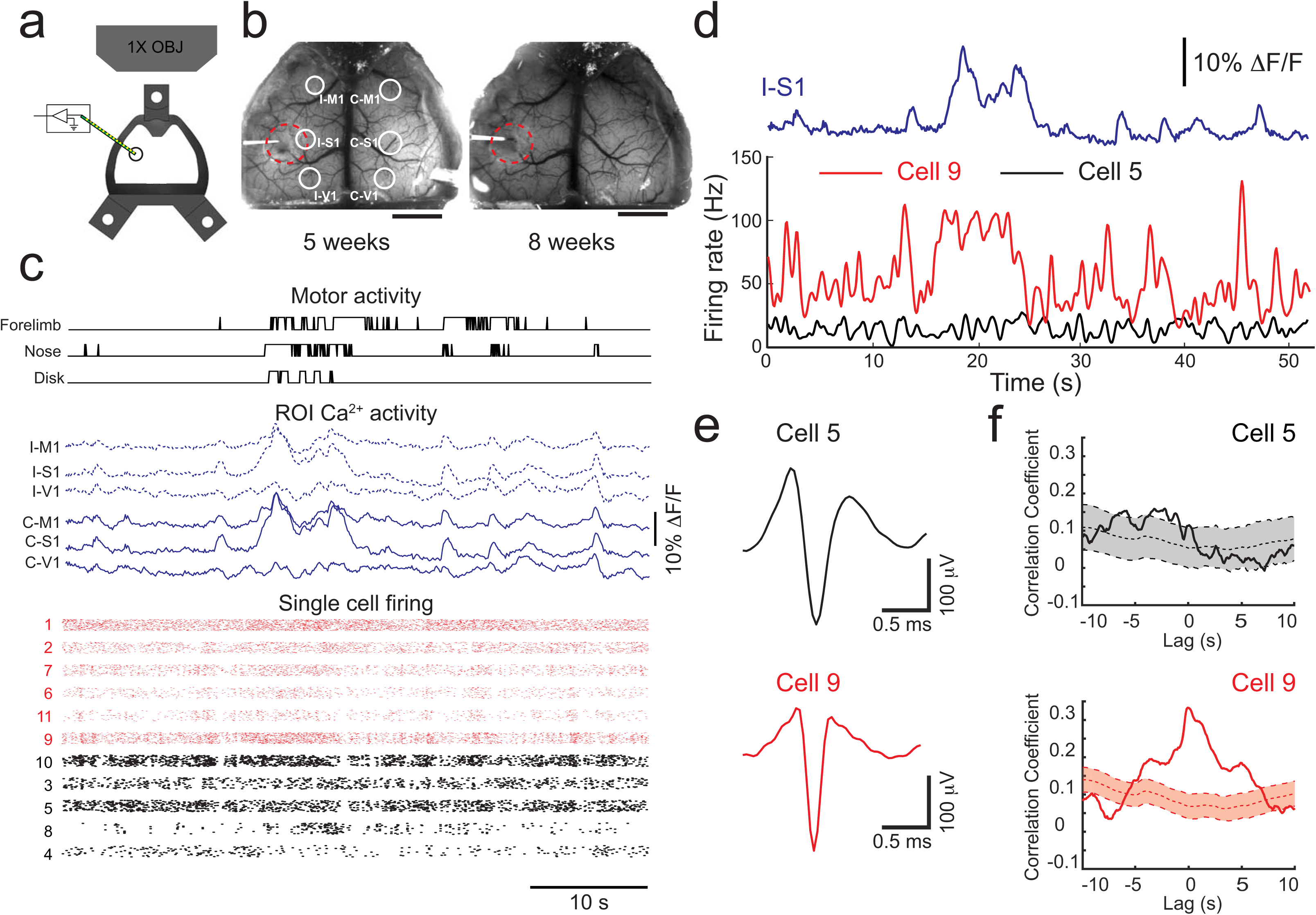
See-Shells can be modified to allow simultaneous extracellular recordings with wide-field Ca^2+^ imaging and behavioral tracking. **(a)** Schematic of implanted See-Shells with perforation over the primary somatosensory cortex allowing insertion of a multi-channel silicon-based neural probe. **(b)** Photographs of a Thy1-GCaMP6f mouse implanted with a perforated See-Shell taken during two experimental sessions. Red dashed lines indicate the outline of perforation. White circles indicate regions of interest (ROIs) analyzed for extracting Ca^2+^ traces. I-M1: ipsilateral primary motor cortex, I-S1: ipsilateral primary somatosensory cortex, I-V1: ipsilateral primary visual cortex, C-M1: contralateral primary motor cortex, C-S1: contralateral primary somatosensory cortex, C-V1: contralateral primary visual cortex. Scale bars indicate 2 mm. **(c)** Simultaneously recorded disk, nose, and forelimb movements, aligned with Ca^2+^ activity traces in the ROIs indicated in **(b)**, and single unit spike raster plots recorded from the multi-channel silicon-based neural probe. Red raster plots indicate neurons with spike firing rates correlated with Ca^2+^ activity in I-S1. Individual points from each cell were slightly shifted in a randomized fashion in the y-axis for ease of visualization. **(d)** Spike firing rates of two representative cells, with one that was correlated with Ca^2+^ activity in I-S1 (cell 9) and one that did not show correlation (cell 5). **(e)** Average action potential waveforms of the two cells shown in **(d). (f)** Cross-correlation of firing rates of representative cells in **(d)** and Ca^2+^ activity in I-S1 with 95% confidence interval of cross-correlations with one thousand bootstrapped shuffled trials of the firing rate to determine significance.

Thus, See-Shells allow multi-scale imaging in the mouse cerebral cortex during a wide-range of head-fixed behaviors. The capability to image in the same animal structurally at subcellular resolution as well as Ca^2+^ activity at the cell and mesoscale level will provide new insights between factors such as physical structure and neural state.

### Perforated See-Shells allow multi-modal and bi-directional neural interfacing

Another advantage of the See-Shell design and PET film is incorporation of additional modalities to record and/or perturb neural structures. For example, combining wide-field Ca^2+^ activity monitoring while simultaneously recording neural firing from localized circuits will enable determination of how global cortical activity relates to local circuit activity^23^. See-Shells were engineered with ∼1.5 mm perforations over the primary somatosensory cortex to introduce a 16 channel, silicon-based recording probes (**Fig. 5a**). The perforations were made prior to implantation and sealed with quick setting silicone sealant that could be removed during experiments (n = 3 Thy1-GCaMP6f, **Fig. 5b**).

Mesoscale Ca^2+^ imaging was performed simultaneously with the single cell recordings in left primary somatosensory cortex and high-speed behavioral recording during awake head-fixation (**Fig. 5c**). Ca^2+^ activity from six regions of interest (ROIs) in the bilateral motor (M1), somatosensory (S1), and visual cortices (V1) show robust co-activation of multiple homotopic regions. Single cell electrophysiology recordings revealed a subset of neurons with increased firing rates correlated with mesoscale Ca^2+^ activity in the ipsilateral primary somatosensory cortex (I-S1) (**Fig. 5c** and **d**). To assess the relation between the individual cell firing and the Ca^2+^ activity, spike firing rate of each neuron was cross-correlated with the Ca^2+^ signals in I-S1. Ca2+ activity was then cross-correlated with 1000 randomly shuffled trials of the spike firing rate of each cell. As shown for two representative cells, one has spike firing correlated with the Ca^2+^ signals in I-S1 (cell 9) and one that did not (cell 5, **Fig. 5e** and **f)**. At zero lag, 6 of the 11 neurons had significant correlation with the Ca^2+^ signals in I-S1 (correlation coefficient > mean + 1.96 s.d. of the bootstrapped traces). These experiments suggest that activities of individual neurons are diverse in terms of their correlation with mesoscale activity. See-shells thus allow us to observe cortical activity at multiple scales and understand their significance to behavior.

Finally, we demonstrate that perforations in See-Shells introduced after chronic implantation can be used to perturb neural circuits with intracortical microstimulation, which has been widely used to assess cortical connectivity and function including effects on downstream targets. In a subset of Thy1-GCaMP6f mice (n = 3, **Fig. 6a**), the PET film was carefully punctured with a sterile syringe needle to introduce microstimulation electrodes. Stimulation resulted in robust activation of both hemispheres in the awake and anesthetized states (**Fig. 6b** and **c**) with significantly prolonged responses in the anesthetized compared to the awake mouse. The full width at half maximum (FWHM) of the post-stimulus Ca^2+^ fluorescence response was significantly longer in all ROIs examined during isoflurane anesthesia (**Fig. 6d,** t-test, p < 0.001). These results with See-Shells in the awake animal extend previous flavoprotein imaging observations in anesthetized mice showing that microstimulation of the primary motor cortex co-activates homotopic regions via the corpus callosum^38^. Therefore, See-Shells can be used to study the effects of perturbing localized regions and suggest that the arousal state alters cortical dynamics. While the present study employed intracortical microstimulation, the methodology could be easily compatible with optogenetic or chemical stimulation of cortical or sub-cortical brain regions.

**Figure 6:**
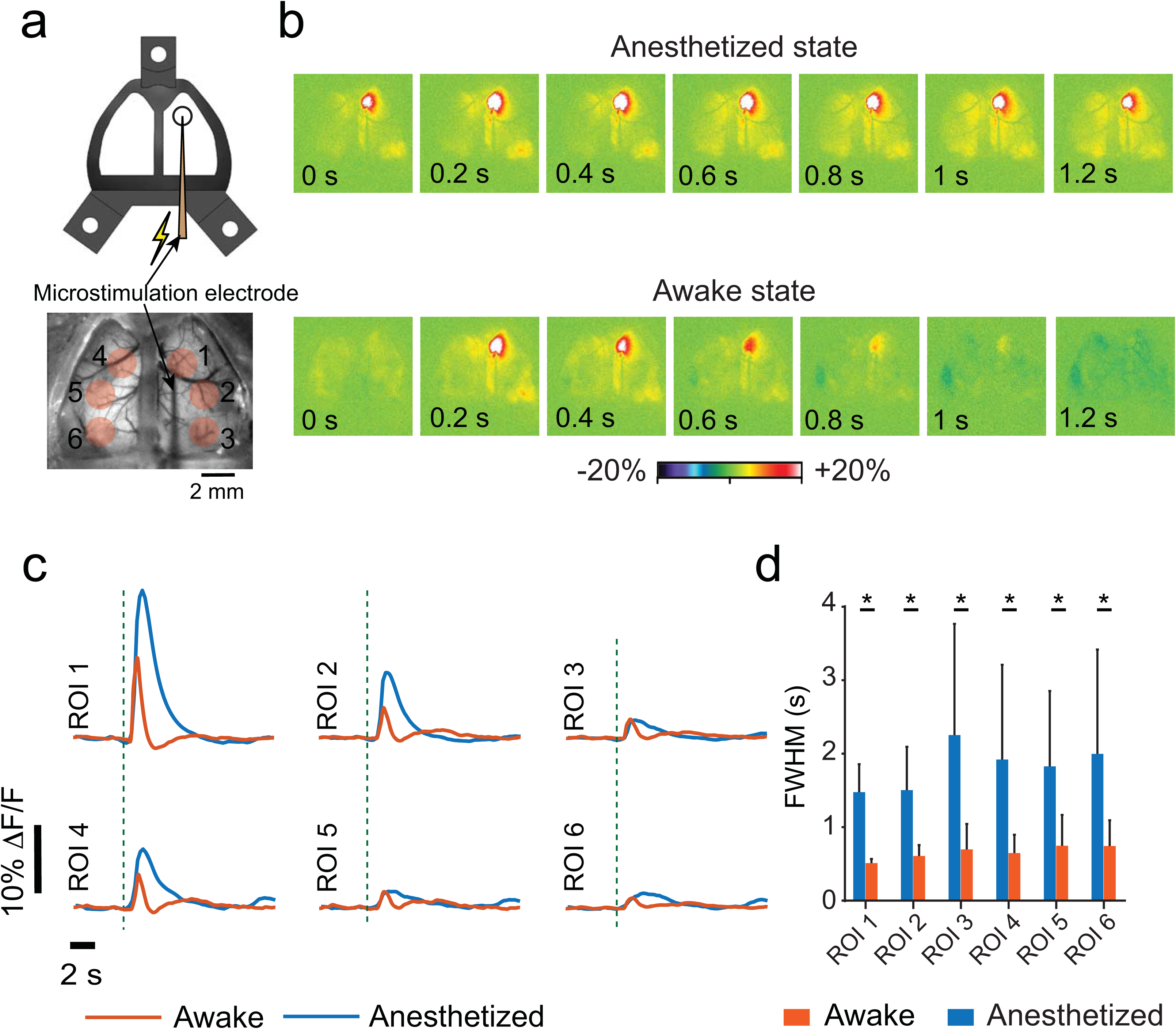
Modified See-Shells allow for cortical microstimulation during wide-field Ca^2+^ imaging. **(a)** *Top:* Cartoon schematic of a microstimulation electrode inserted through a perforated See-Shell. *Bottom:* a wide-field image showing microstimulation electrode inserted at the primary motor cortex. Red circles indicate regions of interest (ROIs) analyzed. **(b)** Pseudocolor plots of normalized change in fluorescent intensity in response to primary motor cortex microstimulation *top:* under isoflurane anesthesia, *bottom:* awake. **(c)** Average normalized Ca^2+^ activity traces in response to microstimulation of primary motor cortex for different ROIs indicated in **(a)**. ROI 1: primary motor cortex and stimulation site; ROI 2: ipsilateral somatosensory cortex; ROI 3: ipsilateral visual cortex; ROI 4: contralateral motor cortex; ROI 5: contralateral somatosensory cortex; and ROI 6: contralateral visual cortex. Dashed lines indicate the time of stimulus. **(d)** Comparison of FWHM in the different ROIs under anesthetized and awake conditions.

## DISCUSSION

We developed See-Shells, transparent, morphologically conformant polymer skulls that allow optical access to a large part of the dorsal cerebral cortex for high-resolution structural and functional imaging. These windows can be implanted for long periods and remain functional for over 250 days. In line with estimates from recent studies using curved glass windows^22^, See-Shells provide optical access to ∼1 million neurons from the cortical surface. In addition, optical imaging with See-Shells can be combined with other modalities. Perforation of the PET film allows access to the brain underneath the implant and here we demonstrated wide-FOV Ca^2+^ imaging with intracortical microstimulation and electrophysiological recordings.

The optical properties of PET compare favorably with glass coverslips when evaluated by 2P or FLIM imaging. The latter open up FLIM intra-vital brain imaging of auto-fluorescence using PET windows^39-41^. For example the intrinsically fluorescent metabolites nicotinamide adenine dinucleotide hydrogen (NADH) and flavin adenine dinucleotide (FAD) are widely used *in vivo* to record label-free cellular activity based on their oxidation state^42-45^. Changes in the lifetime of both coenzymes are used to monitor the biological microenvironment^43^, including in intra-vital studies^46^. Thus, while the current goal is to use PET to realize transparent skulls for cortex-wide imaging, the flexibility, optical clarity and biocompatibility demonstrate the feasibility of engineering anatomically realistic windows for intra-vital imaging in a wide variety of organs such as mammary^47^ and lung^48^.

See-Shells can be implanted using simple modifications to well established chronic cranial window implantation procedures^49,50^, with the major change being the removal of large sections of the skull above the dorsal cerebral cortex. In this study, we utilized a robot that uses surface profiling to guide a computer numerical controlled (CNC) mill to perform the craniotomy^51^. Automation enabled reliable removal of the bone without damage to the underlying dura and brain and also allowed precise positioning of the implant relative to bregma. We also performed the craniotomy manually for See-Shells implantations on *tg/tg* mice, demonstrating that automation of the craniotomy is not a pre-requisite for successful implantation, although it could help investigators quickly adapt these tools for their research.

High quality mesoscopic Ca^2+^ imaging in the awake animal has been performed in mice implanted with See-Shells for 36 weeks, the longest period tested to date. Chronic imaging over this duration provides the opportunity for very long-term studies of brain development, plasticity and learning, disease processes, and evaluation of new therapies. For developmental studies, the primary limitation will be skull growth in the post-natal period. Mouse skull sutures that fuse do so by ∼45 days of age, and most cranial expansion is complete by 6 weeks^52,53^. However, the majority of skull growth is complete earlier (∼ 2-3 weeks of age), suggesting the windows could be implanted in younger animals.

Several aspects of the design and fabrication of the See-Shells are widely adoptable and highly flexible. See-Shells can be fabricated using desktop tools, are inexpensive (< $20 each) and therefore well-within the capabilities of most laboratories. Although the cranial implants were developed for the dorsal cerebral cortex with its fairly regular convex surface, the design can be modified for a variety of skull morphologies. Future versions can be designed to cover not only the dorsal cerebral cortex, but other regions including the olfactory bulb, cerebellum, and more lateral cortical regions such as the auditory cortex.

See-Shells could also be engineered for optically interfacing with complex and mobile anatomical structures such as the spine. The 3D printed frame can be modified to incorporate mounting features to precisely attach miniaturized microscopes^54^ and devices for wirelessly infusing pharmacological agents or performing optogenetic stimulations^55^. Recently, ultra-miniaturized lens-less fluorescence microscopes with thickness less than 1 mm have been developed^56^. Engineering See-Shells embedded with these miniaturized lens-less imaging systems offers the possibility of volumetrically mapping the activity of the whole cortex during freely moving animals.

The ability to simultaneously monitor local microcircuit activity using extracellular recordings combined with wide-FOV Ca^2+^ imaging offers the potential to integrate the contribution of local microcircuits to mesoscale activities. We show that both mesoscale and single unit activities correlated with motor activity. Additional information about the cortical state is likely represented in the complex spatio-temporal patterns of activity observed with mesoscale imaging. See-Shells would allow systematic multi-scale studies tying activity at microcircuits to large scale network activity. Using See-Shells to combine wide-field Ca^2+^ imaging with *in vivo* patch clamping methodologies to record from single^57^ and multiple neurons^58^ will help us better understand the how mesoscale network activity influences sub-threshold membrane potential dynamics in individual neurons.

We have used perforated See-Shells to perform intracortical microstimulation. In addition to stimulating electrodes, this methodology can be used to introduce probes for optogenetic or chemical perturbation of cortical and sub-cortical brain regions. These perturbation strategies will be particularly useful to study how activation or inhibition of localized brain regions or circuits affect global cerebral cortical activity. This includes both specific pathways such as the cerebello-thalamo-cortical^59,60^, basal ganglia-thalamo-cortical projections^61,62^, or the more diffuse neuromodulatory projections such as noradrenergic inputs from the brain stem^63,64^ or cholinergic inputs from the basal forebrain^65^.

With their similar genetics, anatomy, physiology, and behavioral repertoire, non-human primates (NHPs) provide the closest animal model to humans for understanding both normal functions as well as disease^66-68^. Emerging genetic modification techniques, including CRISPR, make generating transgenic NHPs with broad expression of Ca^2+^ reporters a near possibility^69^. Techniques for long-term 2P imaging in NHPs are emerging^70^. The See-Shells methodology can be adapted to build customized implants derived from computed-tomography (CT) scans of the skull, enabling imaging of neural activities across centimeters of the NHP cortex.

While the See-Shells allow sub-cellular resolution imaging across a large FOV, high resolution imaging needs to be done from small regions. Imaging across the whole FOV simultaneously at high resolution is currently not possible. Recently developed wide-field 2P imaging systems^15-17^ allow simultaneous imaging of nearly an entire hemisphere. Extending such optical systems to simultaneously image the whole dorsal cortex at cellular resolution would be very powerful. We have performed wide-FOV mesoscale imaging while simultaneously inserting neural probes for extracellular recordings and microstimulation. It is difficult to do such experiments with 2P microscopes, given the short working distance of high numerical aperture and high magnification objectives. In the future, it may be possible to perform random access 2P imaging across the whole FOV using customized long working distance high-resolution objectives, or by using optical relays between the PET surface and the objectives to allow more room for introduction of recording probes^71^. Neural probes that are specifically designed to be compatible with 2P imaging could also be engineered, or flexible neural probes that can be reconfigured after implantation to provide unimpeded optical access could be used^72^.

## ACKNOWLEDGEMENTS

SBK acknowledges funds from the Mechanical Engineering department, College of Science and Engineering, MnDRIVE RSAM initiative of the University of Minnesota, McGovern Institute Neurotechnology (MINT) fund, National Institutes of Health (NIH) 1R21NS103098-01. MR was supported by 3R21 NS103098-01S1. LG was supported by the University of Minnesota Informatics Institute (UMII) Graduate Research fellowship. TJE was supported, in part, from NIH grants RO1 NS18338 and R37 NS040389. KWE acknowledges funding from DARPA grant N66001-17-2-4010.

We would like to thank Bonita Van Heel at Minnesota Dental Research Center for Biomaterials and Biomechanics for help with micro-CT scanning experiments. We would also like to acknowledge Dr. Mark Sanders and Jason Mitchell at the UMN University Imaging Center where all 2P intensity imaging experiments were conducted. We also acknowledge Drs. Jenu Chacko and Jayne Squirrell of the UW-Madison who assisted with cell culture and viability experiments for FLIM.

## CONTRIBUTIONS

L.G. and S.B.K. conceptualized the technology. L.G., R.E.C, T.J.E., K.W.E. and S.B.K. designed experiments. L.G., R.E.C., M.L.R., J.D., G.C., J.H., M.A.K.S., L.H., N.M., M.M.G., S.L.W. and S.B.K. conducted the experiments. L.G., R.E.C., G.C., M.A.K.S., A.N., T.J.E. and S.B.K. analyzed the data. L.G., R.E.C., M.A.K.S., T.J.E., K.W.E and S.B.K wrote the paper.

## METHODS

### See-Shells design and fabrication

A rodent stereotaxic instrument (David Kopf Instruments Inc.) was modified to have CNC milling capabilities by incorporating a programmatically controlled 3-axis motorized manipulator (MTS25-Z8, Thorlabs)^51^. A handheld mill (Rampower, Ram Products Inc.) fitted with a 200 μm diameter end mill (Harvey tools Inc.) was mounted on the 3-axis manipulator using a custom adaptor plate. A custom computer program was written in LabVIEW (National Instruments Inc.) to control the position of the manipulator.

All of the animal studies were approved by and conducted in conformity with the Institutional Animal Care and Use Committee of the University of Minnesota. An adult C57BL/6 male mouse (8-week, #000664, Jackson Laboratories) and a *tg/tg* male mouse (16-week) were used for skull surface profiling. In each experiment, the mouse was anesthetized using isoflurane in oxygen (4-5% induction, 0.8-1.5% maintenance). The scalp was shaved and sterilized using standard aseptic procedures, after which the mouse was head-fixed in the stereotaxic instrument. The scalp covering the dorsal skull surface was excised and fascia removed using a micro-curette to prepare the skull for surface profiling.

The end mill mounted on a motorized stage was carefully lowered under visualization using a stereo-zoom microscope (M80, Leica) until the end mill tip made contact with the skull surface at bregma and the coordinates were registered in the LabVIEW program. This served as the origin of a Cartesian coordinate system. The LabVIEW program then raised the end mill 0.5 mm above bregma and moved it laterally to the first profiling point (**Supplementary Figs. 1a** and **2a**). The end mill was carefully lowered until it made contact with the skull surface and the z-coordinate was registered. The program then raised the end mill by 0.5 mm and moved it laterally to the next profiling point. The process of registering the z-coordinate was repeated at 85 profiling points on the dorsal skull surface of the C57BL/6 mouse and 134 points on the *tg/tg* mouse. These data were used to construct 3D point clouds to define the skull surfaces (**Supplementary Figs. 1b** and **2b**).

The point cloud was imported into a computer-aided design (CAD) software (Solidworks, Dassault Systèmes). Points along the medial-lateral direction were used to define 3D curves and to interpolate a 3D surface mimicking the skull surface (**Supplementary Figs. 1c** and **2c**). This 3D surface was then extruded to 0.6 - 0.8 mm thickness to create a solid surface which was then used as a template for defining the structural frame of the See-Shell (**Fig. 1a, Supplementary Figs. 1d** and **2d**). The CAD files used for 3D printing of the frame are available for download (**Supplementary File 1**).

See-Shells were fabricated in a multistep process illustrated in detail in **Supplementary Figure 3**. First, the See-Shell structural frame was 3D-printed out of polymethylmethacrylate (PMMA) using a desktop stereolithography (SLA) printer (Form 2, Formlabs Inc., **Supplementary Fig. 3a** and **b**). The three holes in the frame were tapped using a 0-80 hand tap (# 15J611, Grainger). A desktop laser jet printer (HP210w, Hewlett Packard Inc.) was used to print an outline matching the See-Shell frame on a 50 µm thick PET film (MELINEX 462, Dupont Inc.). The PET film was cleaned using ethanol and low-lint cleaning tissue (KimWipes, Kimtech Inc.). A pair of scissors was used to cut the PET film using the printed outline as a reference. (**Supplementary Fig. 3c**). This PET film was then aligned to the PMMA frame and bonded using a clear, two-part quick setting epoxy adhesive (Scotch-Weld(tm), 3M Inc.) (**Supplementary Fig. 3d** and **e**). A head-plate was fabricated from a 0.016” sheet of titanium using a water jet cutter (Omax Inc., see **Supplementary File 1** for CAD drawing). The titanium head-plate was designed such that a 25x objective, with 3 mm long working distance could access the whole FOV provided by the See-Shells while also providing adequate mechanical support for the head-fixed experiments.

### PET optical characterization

An inverted 2P laser scanning microscope was used to image 10 µm YG beads (Polysciences Inc., Warrington, PA, cat#18142-2) through three PET film samples and compared to imaging through a #1.5 glass bottom dish. Time domain fluorescence-lifetime imaging (FLIM) was performed using time-correlated single-photon counting (TCSCP) with Becker and Hickl SPC-150 board to determine the fluorescent lifetime of YG beads imaged through PET. An 80 MHz Ti:Sapphire laser (Spectra Physics; Maitai) tuned to the wavelength of 890 nm was used as the excitation source. The excitation and emission were coupled through an inverted microscope (Nikon; Eclipse TE300) with a 20x air immersion objective (Nikon, Plan Fluor, N.A. 0.75). A 520/30 nm band-pass emission filter (Semrock, Rochester NY) was also used to selectively collect YG beads fluorescence. FLIM images were collected at 256 x 256 pixel resolution with 30 s acquisition using SPC-830 Photon Counting Electronics (Becker & Hickl GbmH, Berlin, Germany) and Hamamatsu H742MP-40 GaAsP photomultiplier tube (Hamamatsu Photonics, Bridgewater, NJ). To compare light intensity attenuation of PET with glass, the laser power was kept at a fixed value using the calibrated power control on a custom-built 2P microscope and the gain on the photomultiplier tube (PMT) was recorded to reach saturation. Urea crystals were used to determine the Instrumentation Response Function (IRF) with a 445/20 bandpass emission filter (Semrock, Rochester. NY). SPCImage software (Becker & Hickl GbmH, Berlin, Germany) was used to analyze the fluorescence lifetime decay curves. The lifetime decay of each pixel was fit with a single exponential decay which resulted in a *x*^2^ error of 1.05 ± 0.12 (mean ± s.d.). Image analyses to estimate FWHM of fluorescent intensities were performed in Fiji^73^.

### *In vivo* surgical implantation

The procedure for implantation of the See-Shells was adapted from previously reported chronic glass window implantation protocols ^50^. Mice were anesthetized using 1-3% isoflurane in pure oxygen (0.6 mL/min). The scalp was shaved and cleaned using standard aseptic surgical procedures. Eyes were covered with ophthalmic eye ointment (Puralube, Dechra Veterinary Products). Buprenorphine (0.1 mg/kg) and Meloxicam (1-2 mg/kg) were administered subcutaneously for analgesia and managing inflammation, respectively. Mice were then head-fixed using ear bars and a nose cone in the stereotaxic instrument equipped with the CNC milling machine. A feedback regulated heating pad was used to maintain the body temperature at 37^°^C throughout the procedure. Anesthetic depth was assessed every 15 minutes via toe pinch stimulus and anesthesia dosage adjusted as needed. Warm lactated Ringer’s solution was administered subcutaneously every two hours to prevent dehydration. A local anesthetic (1% Lidocaine) was administered subcutaneously at the incision sites. The scalp was then removed to expose the skull covering the dorsal and cerebellar cortices. A micro curette was used to scrape the skull surface to remove fascia.

The CNC milling machine incorporated in the stereotaxic instrument was used to perform automated craniotomies in C57BL/6, Thy1-YFP and Thy1-GCaMP6f mice along a predefined path slightly smaller than the perimeter of the See-Shell (**Supplementary Fig. 1a**) as described previously ^51^. The milling was stopped at depths of 107.85 ± 16.79 µm in C57BL/6 (n = 7 mice), 57.14 ± 16.25 µm in Thy1-GCaMP6f (#024276, Jackson Laboratories) (n = 14 mice), and 70 µm in three Thy1-YFP mice (#003709, Jackson Laboratories). Craniotomies in *tg/tg* mice were performed manually. Prior to removing the milled skull, one or two self-tapping screws (F000CE094, Morris Precision Screws and Parts) were implanted 2-3 mm posterior and 3 mm mediolateral to lambda to assist in anchoring the See-Shell to the skull. The skull was removed using surgical forceps taking care to ensure the dura was intact over the entire exposed brain. The brain was covered with sterilized surgical gauze pad soaked in 0.9% saline. The See-Shell was sterilized by soaking in 70% ethanol for 2-3 minutes followed by rinsing in sterile saline. The gauze pad was removed and the See-Shell was gently placed on top the exposed brain and aligned to the craniotomy. The edge of the See-Shell was attached to the skull around the craniotomy by applying a few drops of cyanoacrylate glue (VetBond, 3M Inc.) using a 29-gauge syringe needle. Dental cement (S380, C&B Metabond, Parkell Inc.) was applied to the periphery of the See-Shell to cement it to the skull surface. Care was taken to ensure the screw holes for fastening the titanium head-plates were not filled in with the uncured dental cement. The dental cement was allowed to fully cure. The titanium head-plate was attached to the frame using 3/32” flat head 0-80 screws on the day of implantation. This was followed by a second round of dental cement application to ensure that three fastening locations were fully enclosed in the cement to make it a structurally rigid implant. To protect the implant and underlying brain from light and physical impacts, an opaque 3D-printed PMMA cap was fastened to the titanium head-plate using 3/16” flat head 0-80 screws.

In a subset of Thy1-GCaMP6f mice implanted (n = 3), the See-Shells had ∼1.5 mm diameter perforations above the primary somatosensory cortex (centered -0.76 mm, -2.47mm AL to bregma). The perforation was sealed using quick setting silicone sealant (KWIK-SIL, World Precision Instruments) on the day of the surgery. The brain could be accessed in multiple experimental sessions across weeks by removing and replacing the silicone seal.

After implantation, mice were allowed to recover on a heating pad until ambulatory and then returned to a clean home cage. All mice were administered Buprenorphine and Meloxicam post-operatively on the day of the surgery as well as the 3 succeeding days to assist with full recovery.

### 2P Imaging

All mice were allowed to recover from surgery for 7 or more days before imaging experiments were attempted. A 2P microscope (Leica SP5II) with a 25x (0.95NA) water immersion objective was used for high-resolution imaging experiments *in vivo*. A Mai: Tai Deepsee (Spectra-Physics) laser tuned to 940 nm wavelength for excitation. Mice were head-fixed under the 2P microscope in a custom designed disk treadmill (**Fig. 4b**). Locations in the FOV were targeted at random locations as illustrated in **Figures 3a** and **4a**.

For structural imaging in Thy1-YFP mice, each z-stack had a FOV of 365 μm x 365 μm (512×512 pixels), starting ∼100 μm above the top surface of the PET film, with images acquired every 2 μm down to 800 μm below the starting plane. In two instance, z-stacks were acquired from 5 adjacent tiles by moving the objective in the medial-lateral direction by 340 µm such that one edge has an overlap of 25 µm.

Ca^2+^ imaging was performed using the same setup in fully awake mice. Z-stacks were acquired every 10 µm from adjacent 365 µm x 365 µm tiles with a 15 µm overlap along one edge. Maximum intensity projects of these z-stacks were constructed and macroscopic feature in these projects were used to determine their coordinates in the corresponding wide-field epifluorescence image. Time series were acquired at 20 Hz (256 x 256 pixels) for 5 minutes at one plane in each tile at depths ranging between 200-300µm.

### Intracortical microstimulation during wide-field imaging

Mice were head-fixed in the custom designed disk treadmill placed under a stereo-zoom microscope under light (0.5-1%) isoflurane anesthesia. A feedback regulated heating pad was used to regulate the body temperature at 37^°^ C. The PET film was carefully perforated (+1 mm, +1 mm, AL to bregma) using a 29-gauge syringe for introduction of an intracortical stimulation electrode. The treadmill was then placed under an epifluorescence microscope (QUANTEM: 5125C, Nikon). A 250 µm diameter tungsten micro-electrode (Lot # 217037, FHC) was introduced into the brain at an angle of 45^°^ (anterior-posterior direction) using a micromanipulator. Imaging was performed using a 1x objective when the animal was anesthetized (0.5-1% isoflurane). Each trial lasted a total duration of 2 minutes sampled at 20 Hz using Metamorph (Molecular Devices Inc.). Stimulation train of 20 pulses (200 µA at 100 Hz) was delivered to the primary motor cortex ∼5 seconds after initiation of each trial. The anesthesia was turned off and mouse was allowed to recover for 1 hour before recording during awake state. Behavior was recorded during awake trials using a high-speed camera (FL3-U3-13Y3M-C, FLIR Inc.) at 20 frames/second to monitor whisker or limb movements.

### Simultaneous extracellular recordings and wide-field imaging

Simultaneous extracellular recording and wide-field imaging were performed on Thy1-GCaMP6f mice that had perforations created in the See-Shells prior to implantation. On the day of the experiment, mice were head-fixed in a custom treadmill under a stereo-zoom microscope under light (0.5-1%) isoflurane anesthesia. The silicone seal covering the perforation was carefully removed and the treadmill was placed under the epifluorescence microscope. A 32-channel probe (Neuronexus, A1×32-Edge-5mm-100-177-A32) was mounted on a motorized stage (MPC 385, Sutter Instruments Inc.) and guided to the center of the perforation to touch the dura at ∼2.58 mm lateral and ∼0.76 mm caudal from bregma. Then, the See-Shell was covered with a conductive gel bath of 1% agarose and the ground electrode was placed in a corner of the gel bath. The recording probe was inserted into the brain in 10 µm steps up to a depth of 1 mm from the pial surface into the cortex using a high precision DC motor (MTS25-Z8, Thorlabs) mounted on the Sutter manipulator at a 45^°^ entry angle.

Recordings from the neural probe were first pre-amplified (RA16PA Medusa PreAmps, Tucker Davis Technologies), then transmitted to a second amplifier and digitizer (RZ2 system, Tucker Davis Technologies). Neural data was sampled at 24 kHz, and band passed at 700-5000 Hz to visualize extracellular single units, or between 0-200 Hz to visualize local field potentials. Simultaneous mesoscale optical imaging was performed at 10 Hz. Behavior was recorded during awake trials using a high-speed camera (FL3-U3-13Y3M-C, FLIR Inc.) at 20 frames/second. At the end of the experiment, the perforation was covered with the silicone sealant.

### Data analyses

For wide-field 1P imaging data analyses, six ROIs covering the bilateral motor, somatosensory, and visual cortices were defined. Data analyses were performed in Fiji. Average fluorescent intensity was measured for each ROI in each image. A custom MATLAB script was used to calculate the normalized change in fluorescent intensity over the time series of images. Baseline average fluorescence was obtained by averaging fluorescent intensity over the first 4 seconds of the time series. After normalization, the time series was filtered (2-pole Butterworth low-pass filter: 0.3 Hz)^74^.

Multiphoton Ca^2+^ imaging data were analyzed with Fiji and MATLAB. Briefly, for each time series, the moco (MOtion Corrector) plugin^75^ in Fiji was used to correct for motion artifacts. Maximum intensity and standard deviation images were used to identify cells and place ROIs over each cell in the FOV. For each ROI, the average of the pixel intensity was extracted and imported into a custom MATLAB code. Differential fluorescence intensity (ΔF/F_0_) was calculated for each ROI, where F_0_ was the average pixel value in the ROI over the first 10 seconds.

Behavioral image sequences were imported and analyzed in Fiji. Each image sequence was binned 3 x 3. ROIs were placed over the nose, forelimb, hindlimb, and the disk. Changes in average pixel intensity across the ROIs in sequential frames when there was movement detected, or when there was no movement detected, were given a value of 1 or 0, respectively. This allowed for various types of behavior to be quantified. Walking was classified as changes in all ROIs, grooming was quantified when movement was detected in only the forepaw and nose ROIs.

All extracellular recording data were post-processed using custom MATLAB scripts. The raw voltage traces from multiple channels were filtered using a 150^th^ order finite impulse response (FIR) filter with bounds of 800-5000 Hz. The filtered signals were thresholded to detect action potentials using previously described methods^76^. Cells were sorted using linear discriminant analyses and wavelet decomposition^76-78^. Firing rate for each cell was computed using kernel density estimation and smoothing^79^. To determine the relationship between Ca^2+^ signals and firing rates, we generated one thousand bootstrapped shuffled trials of the spike firing rate of each cell^76^ and computed the cross-correlations with the Ca^2+^ activity in I-S1. The cells were categorize as modulated if their correlation coefficient at zero-lag was greater than mean + 1.96 standard deviation of the bootstrapped trials.

### Histology

A subset of mice (n = 3) were fully anesthetized in 5% isoflurane, and transcardially perfused with phosphate buffered saline (PBS) (CAT# P5493-1L, Sigma Aldrich) followed by 4% paraformaldehyde (PFA, CAT# P6148-500G, Sigma Aldrich). The brains were extracted and stored overnight in 4% PFA for fixation as described previously^80^. The brain was sliced into 50 µm slices and then kept in PBS solution containing 100 mM glycine (50046-50G, Sigma Aldrich) for 30 minutes to quench and excess PFA. This was followed by keeping the slices in PBS solution containing 100 mM glycine and 2% Triton X-100 (93443-100ML, Sigma Aldrich) to permeabilize the tissue. Slices were then kept in PBS solution containing Triton X-100 and blocking agent (Goat Serum, CAT# 927502, Biolegend) for 2 hours after which they were incubated in the solution containing 1:1500 primary antibody-Monoclonal Anti-Glial Fibrillary Acidic Protein (GFAP) antibody produced in mouse (G3893-.2ML, Sigma Aldrich) for 24 hours at 4°C. Slices were washed and incubated in solution containing the secondary antibody conjugated with fluorophores (anti-mouse Alexa 488, ab150117, abcam). Next, the slices were thoroughly washed to clear any excess antibody and mounted on glass slides in VECTASHEILD (H-1200, Vector Labs), a mounting medium containing 4’,6-diamidino-2-phenylindole (DAPI). Mounted slices were imaged under an upright confocal microscope (FV1000 BX2, Olympus FluoView). In each mouse, average fluorescent intensity was measured in FiJi in 7 ROIs (500 µm x 500 µm) distributed in the image, with 4 ROIs at the pial surface and 3 ROIs in layer 2/3. This analysis was repeated in slices from three implanted and three non-surgical control mice.

## SUPPLEMENTARY INFORMATION

**Supplementary Video 1:** Mesoscale imaging of spontaneous changes in fluorescent intensity in the dorsal cortex of a Thy1-GCaMP6f mouse during awake head-fixed behavior

**Supplementary Video 2:** 2P imaging of layer 2/3 neurons in the hindlimb region or the primary motor cortex of a Thy1-GCaMP6f mouse during awake head-fixation. Inset: pseudo-color images of change in fluorescent intensity in 350 µm x 350 µm FOV.

**Supplementary File 1:** File archive of CAD design files of See-Shell components.

## SUPPLEMENTARY FIGUES AND TABLES

**Figure S1:**
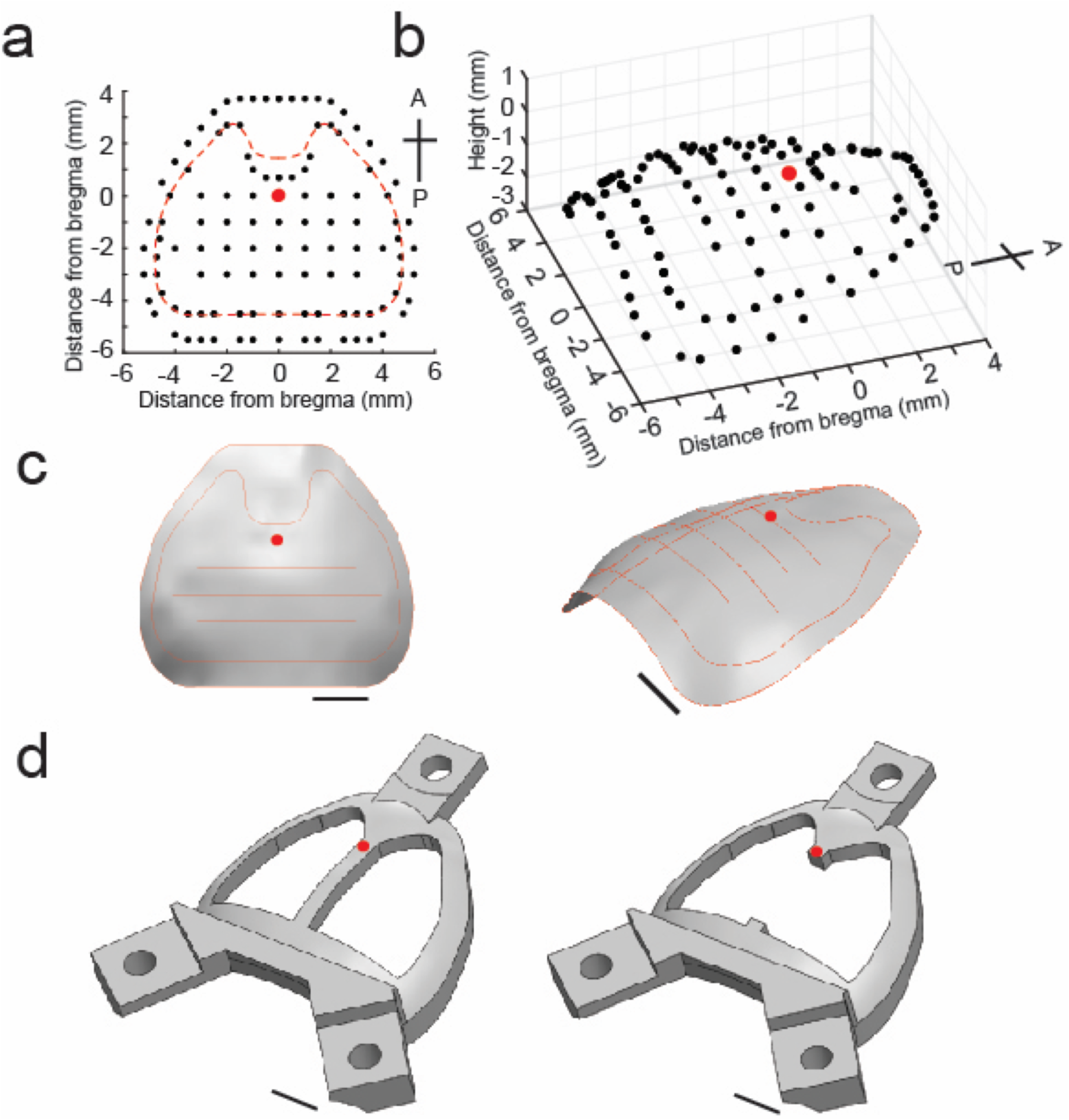
See-Shells designed for C57BL/6 mice: **(a)** Lateral coordinates of 85 points profiled on the dorsal skull surface of a C57BL/6 mouse using the CNC mill in the stereotax (See **Methods**). The dotted line indicates predefined milling path used by the CNC mill during skull excision. **(b)** 3D point cloud generated from surface profiling at the points indicated in **(a)**. **(c)** 3D surface interpolated from the 3D point cloud shown in **(b). (d)** 3D CAD model of the See-Shell’s PMMA frame. Two different versions were used in the experiments, with the version shown on the right used for 2P imaging. Red dots indicate the location of bregma. Scale bars in **c** and **d** indicate 2 mm.

**Figure S2:**
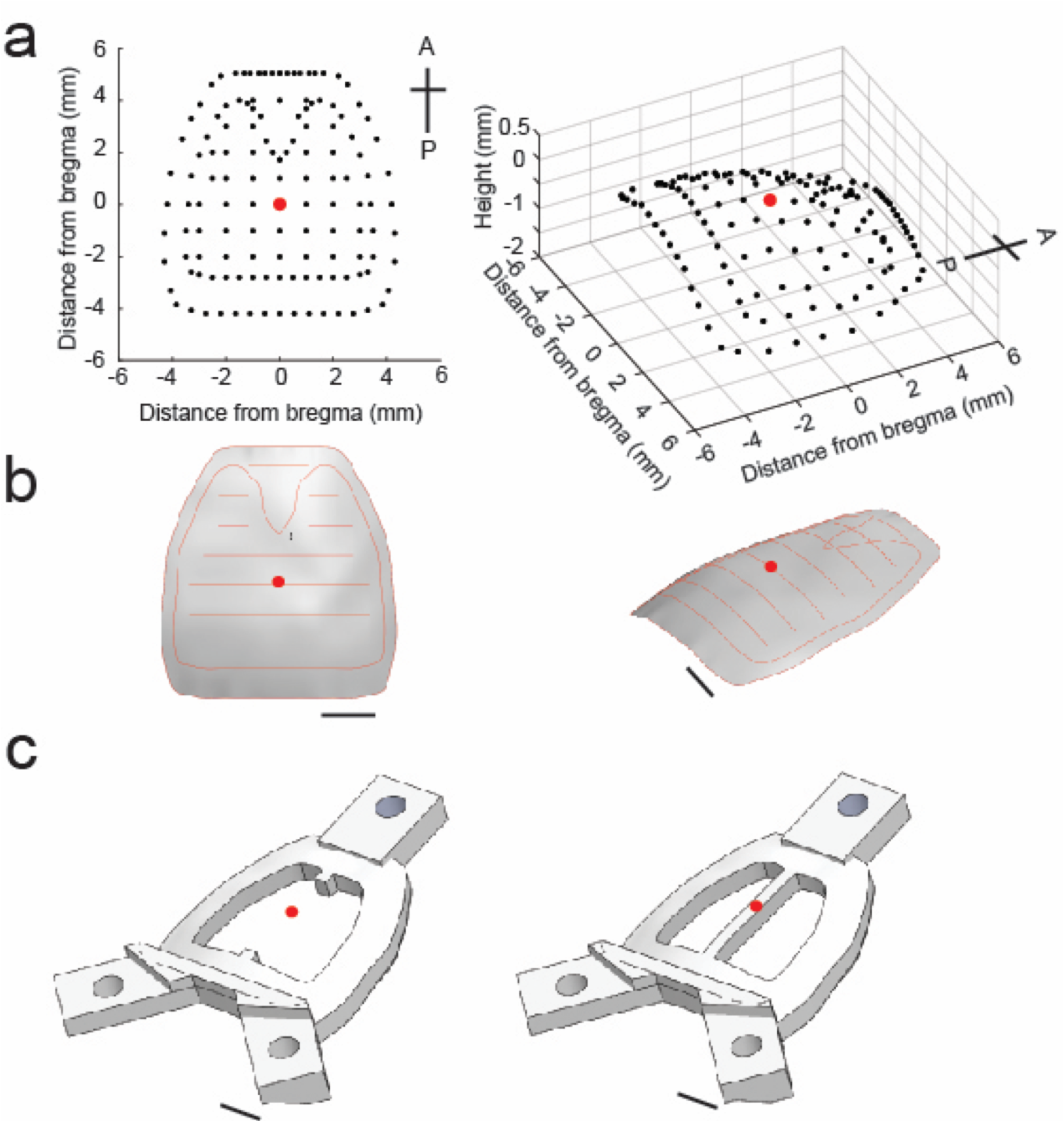
See-Shells designed for *tg/tg* mice: **(a)** Lateral coordinates of 134 points profiled on the dorsal skull surface of a *tg/tg* mouse using the CNC mill in the stereotax (See **Methods**). The dotted line indicates predefined milling path used by the CNC mill during skull excision. **(b)** 3D point cloud generated from surface profiling at the points indicated in **(a)**. **(c)** 3D surface interpolated from the 3D point cloud shown in **(b). (d)** 3D CAD model of the See-Shell’s PMMA frame. Two different versions were used in the experiments, with the version shown on the right used for 2P imaging. Red dots indicate the location of bregma. Scale bars **c** and **d** indicate 2 mm.

**Figure S3:**
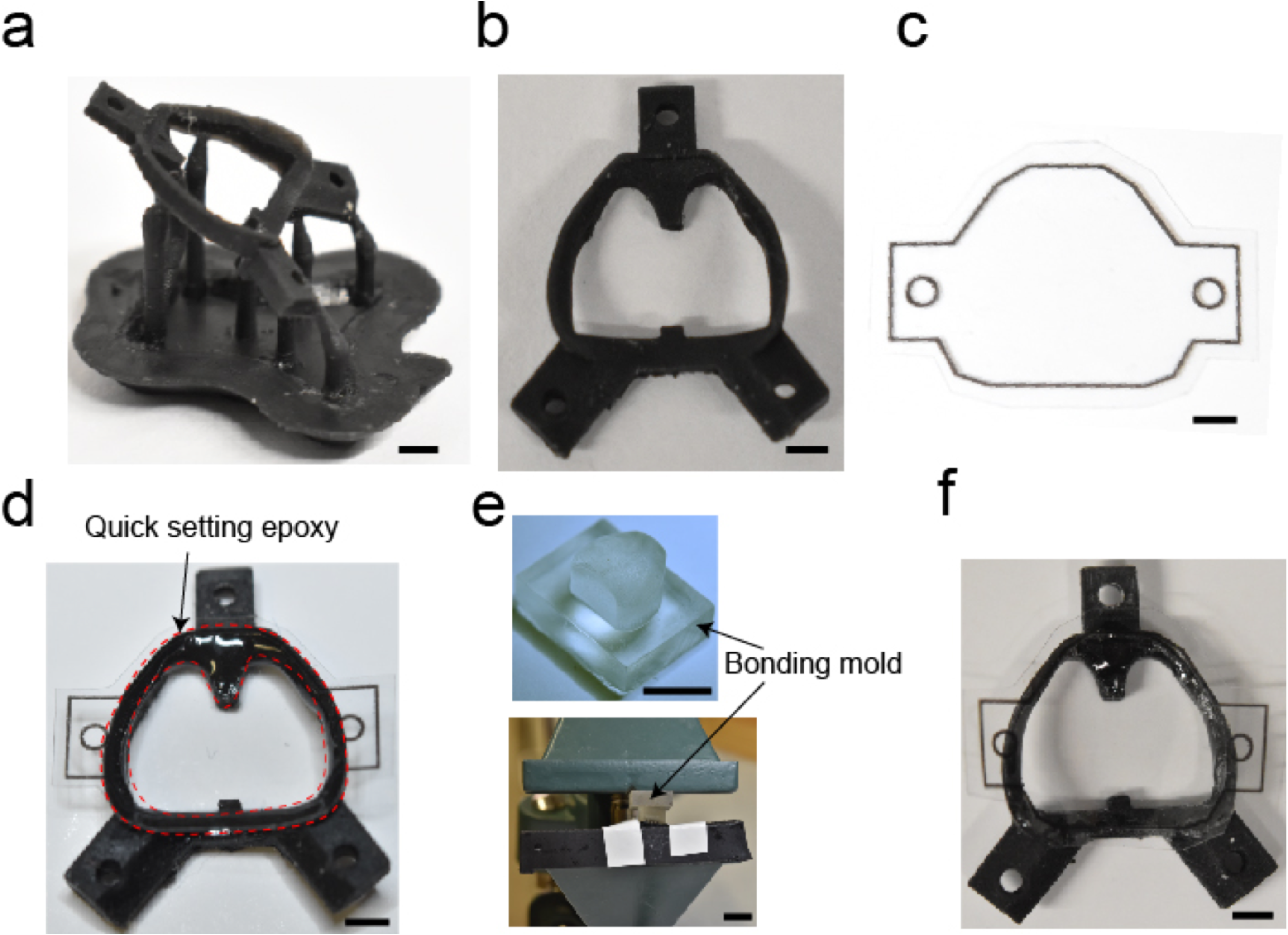
See-Shell assembly: 3D-printed PMMA frame of the See-Shell after retrieval from a 3D printer. **(b)** The frame is cut from support scaffold and washed with %100 Isopropyl alcohol to remove uncured resin. Then the three holes are tapped using a #0-80 tap. **(c)** An outline matching the PMMA frame is printed on a PET film using a desktop printer and is used as a guideline to cut the PET film. **(d)** Bottom surface of the PMMA frame is coated with quick setting epoxy and the cut PET film is aligned with the frame. Red dashed lines delineate areas of epoxy application. **(e)** PET is bonded to the PMMA frame by clamping it down using a custom designed bonding mold in a benchtop vise for 5-10 minutes. **(f)** The implant is removed from the clamp and the excess PET film surrounding the implant cut using a razor blade to realize final implant shown in **Figure 1**. Scale bars in **a, b, c, d** and **f** indicate 2 mm. Scale bar in **e** indicates 1 cm.

**Figure S4:**
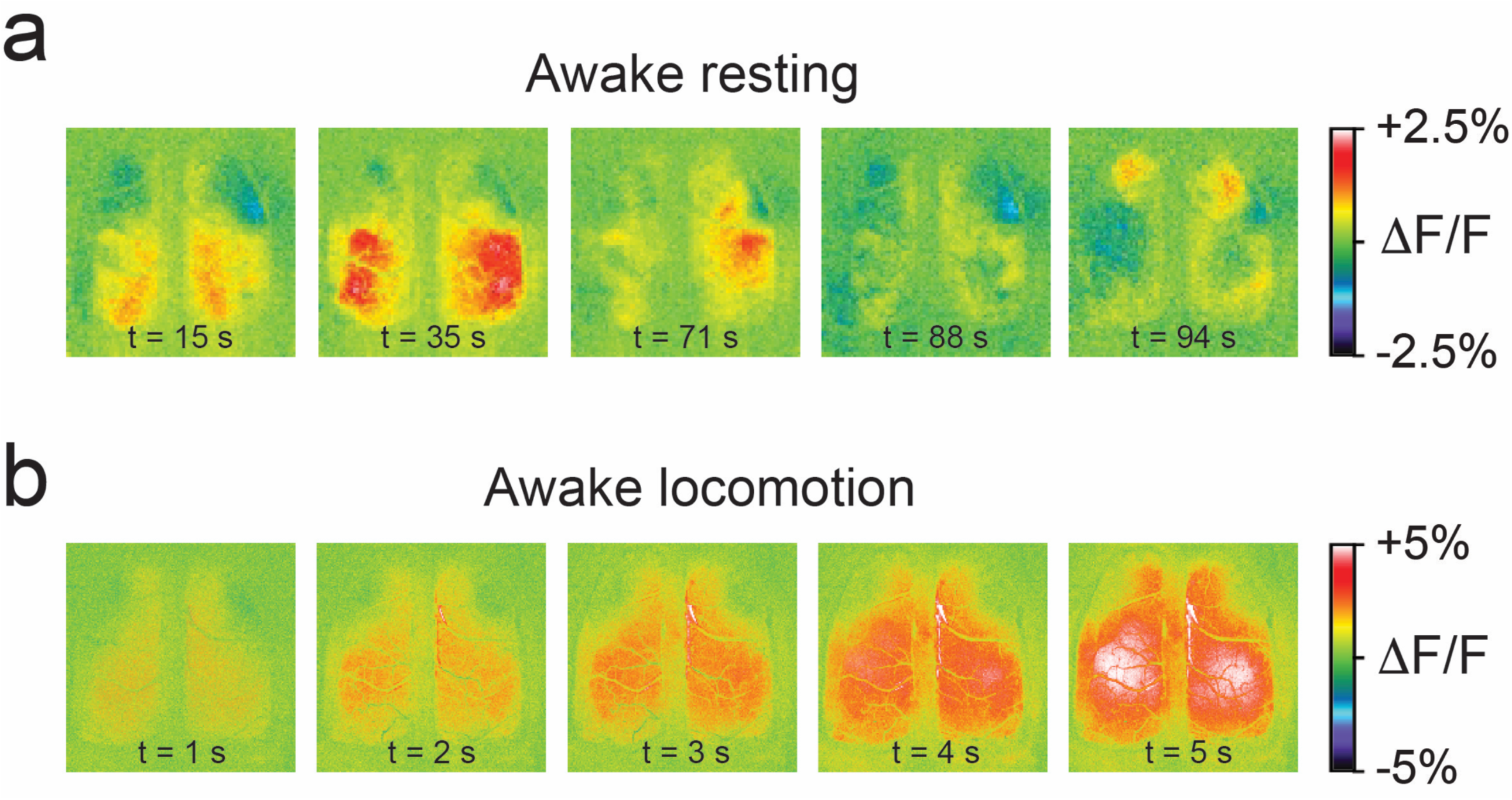
Widely distributed cortical activity during awake behavior. **(a)** Complex mesoscale Ca2+ dynamics, visualized as changes in fluorescence (ΔF/F) during a period of awake resting. **(b)** Mesoscale Ca2+ dynamics visualized as changes in fluorescence (ΔF/F) during a period of locomotion.

**Figure S5:**
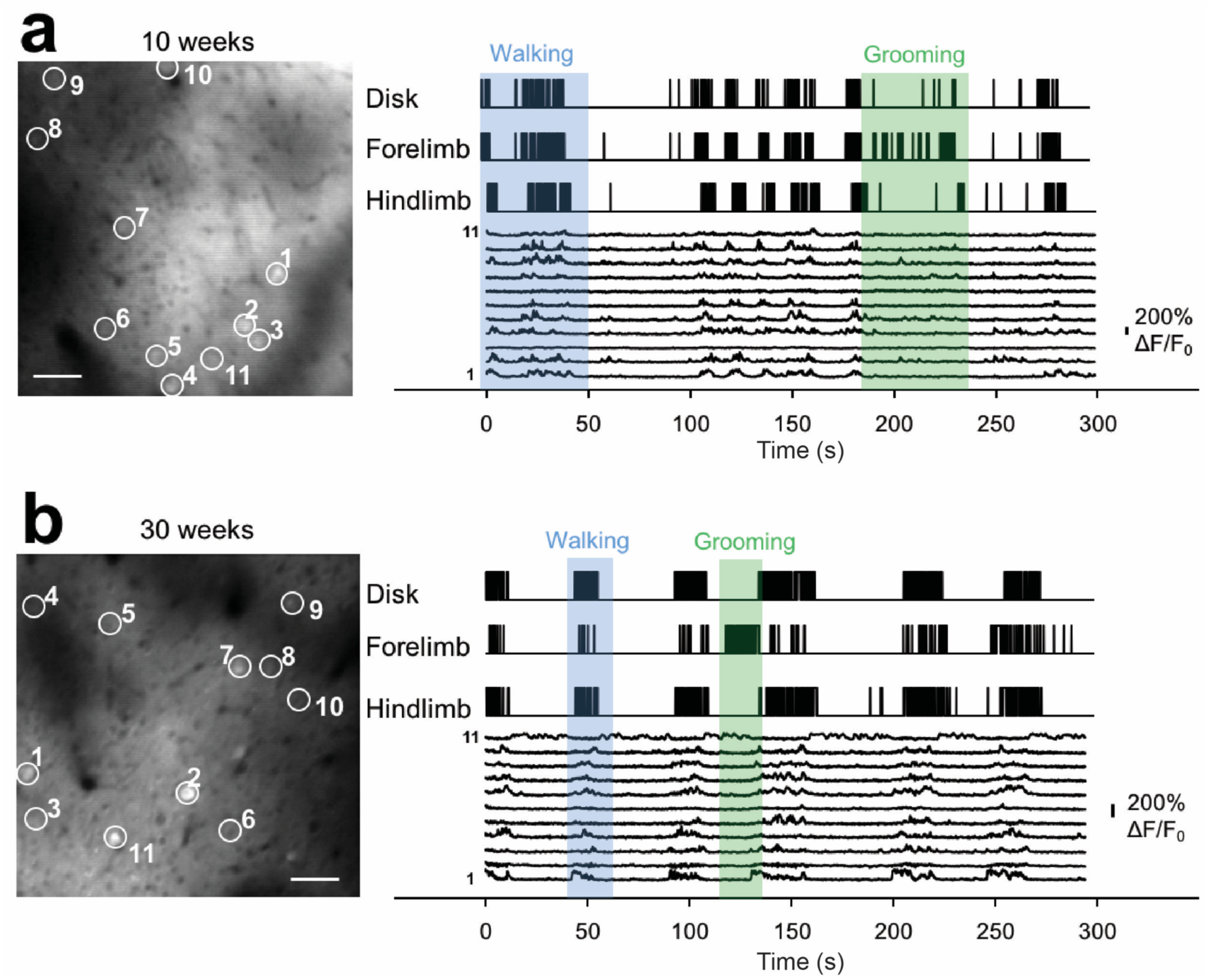
Chronic 2P imaging and behavioral monitoring. **(a)** *Left* Example 2P image from layer 2/3 in the primary motor cortex of a Thy1-GCaMP6f mouse 10 weeks after See-Shell implantation. Individual select neurons (1-11) are outlined by open circles, to show Ca^2+^ transients. Scale bar indicates 50 μm. *Right* Monitoring changes in disk, forelimb, and hindlimb positions with high-speed cameras shows various behaviors such as walking (blue shaded area) and grooming (green shaded area). Single cell Ca^2+^ activity from the select neurons in the 2P image tend to show transients during periods of walking, but not grooming. **(b)** *Left* Example 2P image from layer 2/3 in the primary motor cortex of the same Thy1-GCaMP6f mouse 30 weeks after See-Shell implantation. Optical clarity is still sufficient to see individual neurons (1-11). Scale bar indicates 50 μm. *Right* Similar behaviors such as walking (blue shaded area) and grooming (green shaded area) can still be observed along with single cell Ca^2+^ activity (select neurons in 2P image) at 30 weeks as seen in imaging sessions at 10 weeks (**a**).

**Table S1.**
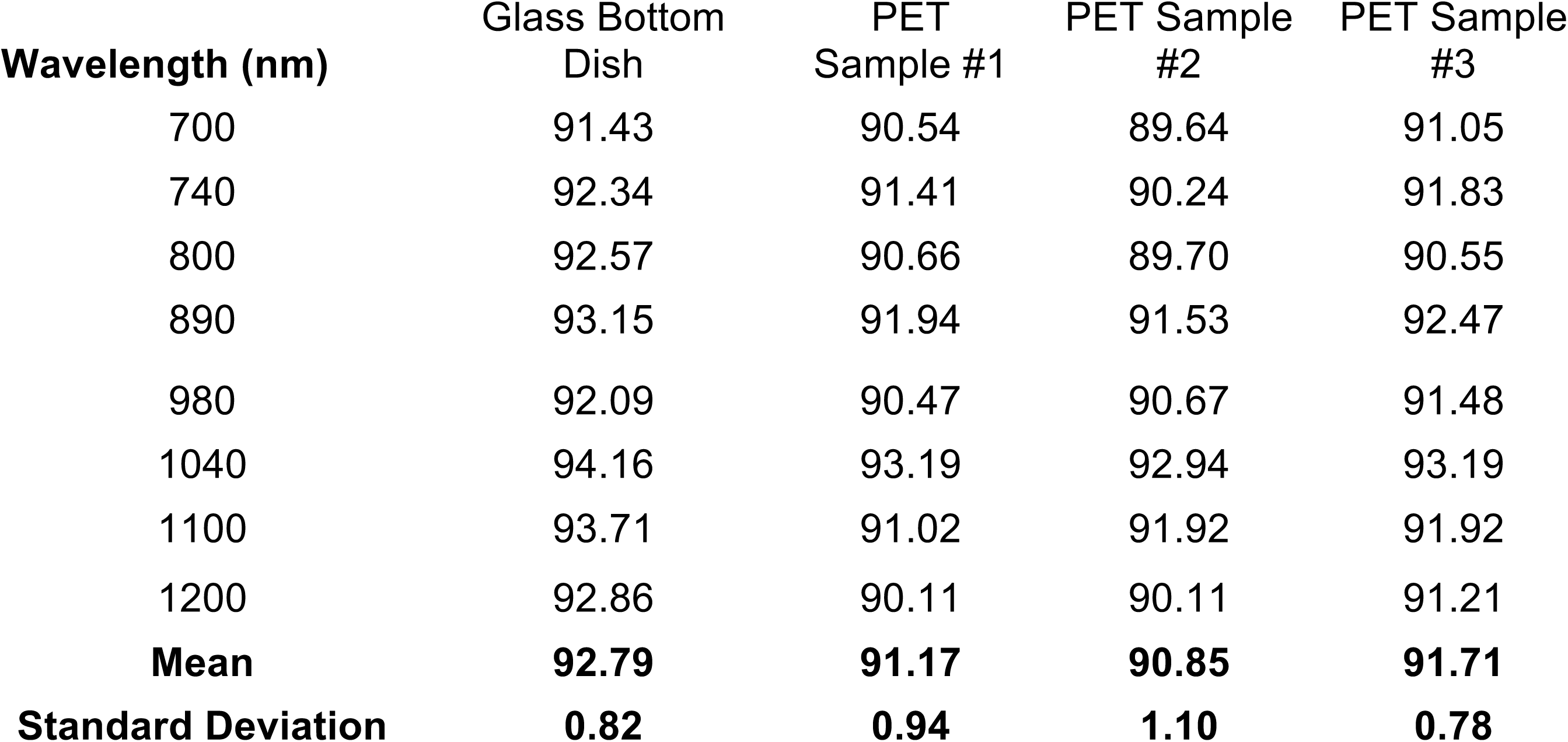
Light Transmission Measurement

## REFERENCES

1. Ferezou, I. et al. Spatiotemporal dynamics of cortical sensorimotor integration in behaving mice. Neuron 56, 907–23 (2007).

2. Dipoppa, M. et al. Vision and Locomotion Shape the Interactions between Neuron Types in Mouse Visual Cortex. Neuron 98, 602–615.e8 (2018).

3. Shimaoka, D., Harris, K. D., Correspondence, M. C. & Carandini, M. Effects of Arousal on Mouse Sensory Cortex Depend on Modality. CellReports 22, 3160–3167 (2018).

4. Denk, W., Strickler, J. H. & Webb, W. W. Two-photon laser scanning fluorescence microscopy. Science 248, 73–6 (1990).

5. Centonze, V. E. & White, J. G. Multiphoton excitation provides optical sections from deeper within scattering specimens than confocal imaging. Biophys. J. 75, 2015–2024 (1998).

6. Abe, T., Maeda, Y. & Iijima, T. Transient increase of the intracellular Ca2+ concentration during chemotactic signal transduction in Dictyostelium discoideum cells. Differentiation. 39, 90–6 (1988).

7. Miyawaki, A. et al. Fluorescent indicators for Ca2+ based on green fluorescent proteins and calmodulin. Nature 388, 882–7 (1997).

8. Nagai, T., Yamada, S., Tominaga, T., Ichikawa, M. & Miyawaki, A. Expanded dynamic range of fluorescent indicators for Ca(2+) by circularly permuted yellow fluorescent proteins. Proc. Natl. Acad. Sci. U. S. A. 101, 10554–9 (2004).

9. Zhao, Y. et al. An Expanded Palette of Genetically Encoded Ca2+ Indicators. Science (80-.). 333, 1888–1891 (2011).

10. Tian, L. et al. Imaging neural activity in worms, flies and mice with improved GCaMP calcium indicators. Nat. Methods 6, 875–81 (2009).

11. Chen, T.-W. et al. Ultrasensitive fluorescent proteins for imaging neuronal activity. Nature 499, 295–300 (2013).

12. Akerboom, J. et al. Genetically encoded calcium indicators for multi-color neural activity imaging and combination with optogenetics. Front. Mol. Neurosci. 6, 2 (2013).

13. Chuong, A. S. et al. Noninvasive optical inhibition with a red-shifted microbial rhodopsin. Nat. Neurosci. 17, 1123–1129 (2014).

14. Klapoetke, N. C. et al. Independent optical excitation of distinct neural populations. Nat. Methods 11, 338–346 (2014).

15. Stirman, J. N., Smith, I. T., Kudenov, M. W. & Smith, S. L. Wide field-of-view, multi-region, two-photon imaging of neuronal activity in the mammalian brain. Nat. Biotechnol. (2016). doi:10.1038/nbt.3594

16. Tsai, P. S. et al. Ultra-large field-of-view two-photon microscopy. Opt. Express 23, 13833–47 (2015).

17. Sofroniew, N. J. et al. A large field of view two-photon mesoscope with subcellular resolution for in vivo imaging. Elife 5, 264–266 (2016).

18. Holtmaat, A. et al. Imaging neocortical neurons through a chronic cranial window. Cold Spring Harb. Protoc. 7, 694–701 (2012).

19. Vanni, M. P. & Murphy, T. H. Mesoscale transcranial spontaneous activity mapping in GCaMP3 transgenic mice reveals extensive reciprocal connections between areas of somatomotor cortex. J. Neurosci. 34, 15931–46 (2014).

20. Silasi, G., Xiao, D., Vanni, M. P., Chen, A. C. N. & Murphy, T. H. Intact skull chronic windows for mesoscopic wide-field imaging in awake mice. J. Neurosci. Methods 267, 141–9 (2016).

21. Drew, P. J. et al. Chronic optical access through a polished and reinforced thinned skull. Nat. Methods 7, 981–984 (2010).

22. Kim, T. H. et al. Long-Term Optical Access to an Estimated One Million Neurons in the Live Mouse Cortex. Cell Rep. 17, 3385–3394 (2016).

23. Xiao, D. et al. Mapping cortical mesoscopic networks of single spiking cortical or sub-cortical neurons. Elife 6, (2017).

24. Faraj, M. G., Ibrahim, K. & Ali, M. K. M. PET as a plastic substrate for the flexible optoelectronic applications. Optoelectron. Adv. Mater. Rapid Commun. 5, 879–882 (2011).

25. Khan, W., Muntimadugu, E., Jaffe, M. & Domb, A. J. Implantable Medical Devices. in Focal Controlled Drug Delivery, Advances in Delivery Science and Technology 33–59 (2014). doi:10.1007/978-1-4614-9434-8

26. Becker, W., Bergmann, A. & Biskup, C. High resolution TCSPC lifetime imaging. Proc. … 4963, 1–10 (2003).

27. Fletcher, C. F. et al. Absence epilepsy in tottering mutant mice is associated with calcium channel defects. Cell 87, 607–617 (1996).

28. Green, M. C. & Sidman, R. L. Tottering-a neuromuscular mutation in the mouse: And its linkage with oligosyndactylism. J. Hered. 53, 233–237 (1962).

29. Gray, N. W., Weimer, R. M., Bureau, I. & Svoboda, K. Rapid redistribution of synaptic PSD-95 in the neocortex in vivo. PLoS Biol. 4, 2065–2075 (2006).

30. Nimmerjahn, A., Kirchhoff, F. & Helmchen, F. Neuroscience: Resting microglial cells are highly dynamic surveillants of brain parenchyma in vivo. Science (80-.). 308, 1314–1318 (2005).

31. Holtmaat, A. & Svoboda, K. Experience-dependent structural synaptic plasticity in the mammalian brain. Nat. Rev. Neurosci. 10, 647–58 (2009).

32. Poirazi and Mel, B.W.P., Impact of active dendrites and structural plasticity on the memory capacity of neural tissue. Neuron 29, 779–96. (2001).

33. Chklovskii, D. B., Mel, B. W. & Svoboda, K. Cortical rewiring and information storage. Nature 431, 782–788 (2004).

34. Yang, G., Pan, F. & Gan, W. B. Stably maintained dendritic spines are associated with lifelong memories. Nature 462, 920–924 (2009).

35. Roberts, T. F., Tschida, K. A., Klein, M. E. & Mooney, R. Rapid spine stabilization and synaptic enhancement at the onset of behavioural learning. Nature 463, 948–952 (2010).

36. Feng, G. et al. Imaging neuronal subsets in transgenic mice expressing multiple spectral variants of GFP. Neuron 28, 41–51 (2000).

37. Dana, H. et al. Thy1-GCaMP6 transgenic mice for neuronal population imaging in vivo. PLoS One 9, (2014).

38. Chen, G. et al. Altered levels of the splicing factor muscleblind modifies cerebral cortical function in mouse models of myotonic dystrophy. Neurobiol. Dis. 112, 35–48 (2018).

39. Papour, A. et al. Optical imaging for brain tissue characterization using relative fluorescence lifetime imaging. J. Biomed. Opt. 18, 60504 (2013).

40. Yaseen, M. a et al. In vivo imaging of cerebral energy metabolism with two-photon fluorescence lifetime microscopy of NADH. Biomed. Opt. Express 4, 307–21 (2013).

41. Tejwani, V. et al. Investigation of the NADH/NAD+ratio in Ralstonia eutropha using the fluorescence reporter protein Peredox. Biochim. Biophys. Acta - Bioenerg. 1858, 86–94 (2017).

42. Bird, D. K. et al. Metabolic mapping of MCF10A human breast cells via multiphoton fluorescence lifetime imaging of the coenzyme NADH. Cancer Res. 65, 8766–8773 (2005).

43. Skala, M. C. et al. In vivo multiphoton microscopy of NADH and FAD redox states, fluorescence lifetimes, and cellular morphology in precancerous epithelia. Proc. Natl. Acad. Sci. 104, 19494–19499 (2007).

44. Shibuki, K. et al. Dynamic imaging of somatosensory cortical activity in the rat visualized by flavoprotein autofluorescence. Journal of Physiology 549, 919–927 (2003).

45. Reinert, K. C. et al. Cellular and metabolic origins of flavoprotein autofluorescence in the cerebellar cortex in vivo. Cerebellum 10, 585–599 (2011).

46. Cheng, S. et al. Flexible endoscope for continuous in vivo multispectral fluorescence lifetime imaging. Opt. Lett. 38, 1515 (2013).

47. Szulczewski, J. M. et al. In Vivo Visualization of Stromal Macrophages via label-free FLIM-based metabolite imaging. Sci. Rep. 6, (2016).

48. Entenberg, D. et al.. In vivo subcellular resolution optical imaging in the lung reveals early metastatic proliferation and motility. IntraVital 4, 1–11 (2015).

49. Roome, C. J. & Kuhn, B. Chronic cranial window with access port for repeated cellular manipulations, drug application, and electrophysiology. Front. Cell. Neurosci. 8, 379 (2014).

50. Holtmaat, A. et al. Long-term, high-resolution imaging in the mouse neocortex through a chronic cranial window. Nat. Protoc. 4, 1128–1144 (2009).

51. Ghanbari, L. et al. Principles of Computer Numerical Controlled Machining Applied to Cranial Microsurgery. bioRxiv 280461 (2018). doi:10.1101/280461

52. Vora, S. R., Camci, E. D. & Cox, T. C. Postnatal ontogeny of the cranial base and craniofacial skeleton in male C57BL/6J mice: A reference standard for quantitative analysis. Front. Physiol. 6, (2016).

53. Bradley, J. P., Levine, J. P., Roth, D. A., McCarthy, J. G. & Longaker, M. T. Studies in cranial suture biology: IV. Temporal sequence of posterior frontal cranial suture fusion in the mouse. Plast. Reconstr. Surg. 98, 1039–1345 (1996).

54. Ghosh, K. K. et al. Miniaturized integration of a fluorescence microscope. Nat. Methods 8, 871–878 (2011).

55. Jeong, J. W. et al. Wireless Optofluidic Systems for Programmable In Vivo Pharmacology and Optogenetics. Cell 162, 662–674 (2015).

56. Adams, J. K. et al. Single-frame 3D fluorescence microscopy with ultraminiature lensless FlatScope. Sci. Adv. 3, (2017).

57. Kodandaramaiah, S. B., Talei Franzesi, G., Chow, B. Y., Boyden, E. S. & Forest, C. R. Automated whole-cell patch-clamp electrophysiology of neurons in vivo. Nat. Methods (2012). doi:10.1038/nMeth.1993

58. Kodandaramaiah, S. B. et al. Multi-neuron intracellular recording in vivo via interacting autopatching robots. Elife 7, (2018).

59. Chaumont, J. et al. Clusters of cerebellar Purkinje cells control their afferent climbing fiber discharge. Proc. Natl. Acad. Sci. 110, 16223–16228 (2013).

60. Evarts, E. V & Thach, W. T. Motor mechanisms of the CNS: cerebrocerebellar interrelations. Annu. Rev. Physiol. 31, 451–498 (1969).

61. Shipp, S. The functional logic of corticostriatal connections. Brain Structure and Function 222, 669–706 (2017).

62. Marchand, W. R. Cortico-basal ganglia circuitry: A review of key research and implications for functional connectivity studies of mood and anxiety disorders. Brain Structure and Function 215, 73–96 (2010).

63. Borodovitsyna, O., Flamini, M. & Chandler, D. Noradrenergic Modulation of Cognition in Health and Disease. Neural Plast. 2017, (2017).

64. Szabadi, E. Functional neuroanatomy of the central noradrenergic system. J. Psychopharmacol. 27, 659–93 (2013).

65. Ballinger, E. C., Ananth, M., Talmage, D. A. & Role, L. W. Basal Forebrain Cholinergic Circuits and Signaling in Cognition and Cognitive Decline. Neuron 91, 1199–1218 (2016).

66. Huang, L., Merson, T. D. & Bourne, J. A. In vivo whole brain, Cellular and molecular imaging in nonhuman primate models of neuropathology. Neuroscience and Biobehavioral Reviews 66, 104–118 (2016).

67. Camus, S., Ko, W. K. D., Pioli, E. & Bezard, E. Why bother using non-human primate models of cognitive disorders in translational research? Neurobiology of Learning and Memory 124, 123–129 (2015).

68. Dettmer, A. M. & Suomi, S. J. Nonhuman primate models of neuropsychiatric disorders: Influences of early rearing, genetics, and epigenetics. ILAR J. 55, 361–370 (2014).

69. Park, J. E. et al. Generation of transgenic marmosets expressing genetically encoded calcium indicators. Sci. Rep. 6, (2016).

70. Li, M., Liu, F., Jiang, H., Lee, T. S. & Tang, S. Long-Term Two-Photon Imaging in Awake Macaque Monkey. Neuron 93, 1049–1057.e3 (2017).

71. Lecoq, J. et al. Visualizing mammalian brain area interactions by dual-axis two-photon calcium imaging. Nat. Neurosci. 17, 1825–1829 (2014).

72. Luan, L. et al. Nanoelectronics enabled chronic multimodal neural platform in a mouse ischemic model. J. Neurosci. Methods 295, 68–76 (2018).

73. Schindelin, J. et al. Fiji: An open-source platform for biological-image analysis. Nature Methods 9, 676–682 (2012).

74. Cai, D. J. et al. A shared neural ensemble links distinct contextual memories encoded close in time. Nature 534, 115–8 (2016).

75. Dubbs, A., Guevara, J., Peterka, D. S. & Yuste, R. moco: Fast Motion Correction for Calcium Imaging. arXiv 10, 1–7 (2015).

76. Quiroga, R. Q., Nadasdy, Z. & Ben-Shaul, Y. Unsupervised spike detection and sorting with wavelets and superparamagnetic clustering. Neural Comput. 16, 1661–1687 (2004).

77. Keshtkaran, M. R. & Yang, Z. Unsupervised spike sorting based on discriminative subspace learning. in *2014 36th Annual International Conference of the IEEE* Engineering in Medicine and Biology Society, EMBC 2014 3784–3788 (2014). doi:10.1109/EMBC.2014.6944447

78. Yang, Z., Zhao, Q. & Liu, W. Spike Feature Extraction Using Informative Samples. NIPS 1–8 (2008).

79. Fujisawa, S., Amarasingham, A., Harrison, M. T. & Buzsáki, G. Behavior-dependent short-term assembly dynamics in the medial prefrontal cortex. Nat. Neurosci. 11, 823–833 (2008).

80. Gage, G. J., Kipke, D. R. & Shain, W. Whole Animal Perfusion Fixation for Rodents. J. Vis. Exp. e3564–e3564 (2012). doi:10.3791/3564

